# Hippocampal Ring Finger Protein 10-dependent signaling supports cognitive flexibility

**DOI:** 10.64898/2026.03.31.715507

**Authors:** Elena Romito, Nicolò Carrano, Ana Ribeiro, Maria Italia, Filippo La Greca, Francesca Genova, Laura D’Andrea, Elisa Zianni, Luisa Ponzoni, Gael Barthet, Stephan C. Collins, Mariaelvina Sala, Nico Mitro, Christophe Mulle, Binnaz Yalcin, Elena Marcello, Fabrizio Gardoni, Monica DiLuca, Diego Scheggia

## Abstract

The ability to flexibly adapt behavior to changing environmental contingencies is a core component of brain function and relies on experience-dependent remodeling of neural circuits. While cognitive flexibility has been primarily attributed to prefrontal–striatal networks, the contribution of hippocampus and their underlying molecular substrates remains less understood. Here, we show that the dorsal hippocampus has a key role in cognitive flexibility. In particular, Ring Finger Protein 10 (RNF10)-mediated signaling, linking activation of synaptic NMDARs to specific transcriptional programs in the dorsal CA1, is necessary for cognitive flexibility. In fact, *in vivo* downregulation, through gene deletion and silencing of RNF10, resulting in impaired long-term synaptic plasticity, suppressed cognitive flexibility. This was reflected in the impaired ability to disengage from previously acquired contextual, visual, and spatial information and to adapt behavior to changed context. Overall, our results identified RNF10 as a key *in vivo* player necessary for the balance between cognitive stability and flexibility.

## INTRODUCTION

The ability to adapt behavior to changing environmental demands is a fundamental property of brain function and a key determinant of brain health across the lifespan. This adaptive capacity relies on activity-dependent remodeling of neural circuits that enables the brain to update previously learned associations and adjust behavior when environmental contingencies change (1). A key manifestation of this adaptive capacity is cognitive flexibility. Common descriptions of the neural basis of cognitive flexibility emphasize the contribution of frontal cortical networks interacting with striatal circuits (2, 3). In both non-human primates and rodents, lesions of the orbitofrontal cortex (OFC) impair reversal learning without affecting other components of cognitive flexibility, such as extradimensional set shifting (4–6). These studies have led to the hypothesis that lateral prefrontal cortices support shifts between abstract perceptual dimensions, whereas the OFC, and its associated corticostriatal loops, are required to update stimulus–outcome associations when reinforcement contingencies change (3). Classical models of cognitive flexibility less commonly include hippocampal contributions. However, accumulating evidence suggests that hippocampal circuits may participate in goal-directed behavior, through interaction with prefrontal regions (7), by updating contextual representations and comparing expected and actual outcomes (8–10). Beyond its established roles in episodic and spatial memory (11, 12), hippocampal population activity dynamically tracks latent state transitions during reversal learning, and disruption of hippocampal function impairs adaptation to contingency shifts (9, 13). These findings position hippocampal circuits as active regulators of context-dependent memory updating guiding behaviors. Such adaptive functions rely on experience-dependent modifications of synaptic strength and structure within hippocampal networks (14).

Experience-dependent modifications of synaptic strength and structure contribute to the stabilization and revision of memory representations within hippocampal circuits (15–17). The persistence of long-lasting synaptic changes requires coordinated transcriptional programs initiated by neuronal activity (15–17). N-methyl-D-aspartate receptors (NMDARs), play a central role in excitation–transcription coupling by linking synaptic calcium influx to activity-dependent gene expression necessary for sustained structural and functional plasticity (17–21). Synapto-nuclear messengers mediate communication between activated synapses and the nucleus (22–25). Ring Finger Protein 10 (RNF10) is a synapto-nuclear protein localized at excitatory synapses and directly binding GluN2A containing NMDA receptors (26, 27). Although RNF10 regulates activity-dependent gene expression required for long-term potentiation and spine remodeling in vitro (26, 27), the behavioral implications of RNF10-mediated synapse-to-nucleus signaling in complex behavior have not been established.

Here, we show that disruption of RNF10 in dorsal CA1 selectively impairs reversal learning, increases perseverative errors, and enhances persistence of contextual memory while sparing initial acquisition. At the cellular level, RNF10 loss alters dendritic spine morphology, increases intrinsic excitability, impairs long-term synaptic plasticity, and dysregulates GluN2A–RasGRF2 signaling. These findings identify RNF10-dependent synapse-to-nucleus signaling as a molecular mechanism contributing to hippocampal memory updating and to the balance between cognitive stability and flexibility.

## MATERIALS AND METHODS

### Animals

All procedures involving animals were approved by the local animal use committee and the Italian Ministry of Health (permits #191/2016; #385/2019; #374/2020; #5247B.N.YCK) and were conducted following the Guide for the Care and Use of Laboratory Animals of the National Institutes of Health and the European Community Council Directives. Routine veterinary care and animal maintenance were provided by dedicated and trained personnel. We used 2-to-6-month-old male C57BL/6N mice lacking the RNF10 gene (RNF10 KO) or WT littermates. The C57BL/6N-Rnf10tm1b(KOMP)Wtsi mouse model was obtained from the Knockout Mouse Project (KOMP) Repository (UC Davis). No published studies have been reported on this mouse model. Two to four animals were housed per cage in a climate-controlled facility (ambient temperature of 22 ± 2 °C and relative humidity of 45–65%; 12-h light/dark cycle (lights on at 8:00 am; dark at 8:00 pm) with *ad libitum* access to food and water. Animal handling and surgical procedures were carried out with care taken to minimize discomfort and pain following the ethical guidelines and regulations of the European Parliament and of the council on the protection of animals used for scientific purposes (Directive of 22 September 2010, 2010/63/EU). Experiments were run during the light phase (10:00 am to 5:00 pm). All mice were handled on alternate days during the week preceding the first behavioral tests. Distinct cohorts of mice were used for each experimental approach (behavioral tasks, brain anatomy, spine morphology, and biochemistry).

### Viral vectors

For shRNA experiments, sequences for mouse RNF10 shRNA (mature antisense TCAGGTTGATCTTCTTAGGG) and scramble shRNA (purchased from Origene, Rockville, MD) were subcloned downstream of U6 in the U6-CamKIIa.mCherry-WPRE backbone (provided by Prof. Daniela Mauceri, University of Heidelberg, DE) using BamHI and HindIII restriction enzymes (New England Biolabs, USA). Viral particles (AAV2-U6.scr-CamKIIa.mCherry and AAV2-U6.shRnf10-CamKIIa.mCherry) were prepared by the Viral Vector Facility (VVF) of the Neuroscience Center Zurich (ZNZ). For the rescue experiments, RNF10-ShResistant sequence was obtained by inserting silent point mutations in the RNF10 sequence to avoid ShRNA targeting. The sequence was subcloned downstream of SYN1 promoter. AAV1-SYN1>Myc/Myc/Myc/RNF10shResistant was produced by VectorBuilder.

### Surgical procedures

C57/BL6N male mice were anesthetized with a mix of isoflurane and oxygen by inhalation and mounted into a stereotaxic frame (Kopf Instruments) linked to a digital micromanipulator. Brain coordinates of bilateral injection of AAV in the dorsal CA1 were chosen following the mouse brain atlas: anterior–posterior (AP), −2.00 mm; medial–lateral (ML): ±1.5 mm; and dorsal–ventral (DV): −1.3 mm. AAVs were infused (700 nL) through a 10-μL Hamilton syringe using a microinjection pump at a flow rate of 1.0 µL/min. For the rescue experiments, a 1:1 mixture of AAV1-shRNF10 and AAV1-RNF10-shResistant was prepared immediately before injection. For injections involving RNF10 shRNA or scramble ShRNA alone, the volume of AAV1-RNF10-shResistant was replaced with an equal volume of PBS.

The surgical procedure lasted approximately 20 min for each animal. Mice received carprofen (5 mg/kg) in drinking water for three consecutive days.

## Behavioral tests

### Morris water maze

The Morris water maze test was used to analyze changes in the learning and memory abilities of the mice according to (28) (adapted for mice). A circular water maze (120 cm in diameter × 50 cm in height) was used. A circular hidden platform with a diameter of 10 cm was placed inside the maze, and its surface was maintained at 0.5 cm below the surface of the water. Floating plastic particles were placed on the surface of the water to hide the platform from sight, according to (29). For the habituation trials, the mice were placed in a random area inside the maze and allowed to swim for 60 s. For the acquisition trials, the mice were submitted to four trials per day (with 60-min inter-trial intervals) for four consecutive days, during which each mouse was released into the pool at different starting points and trained to locate a constant platform position. A probe test was performed 24 h after the last trial, during which the platform was removed. Two days later, a reversal task was performed to assess cognitive flexibility. The platform was placed in the opposite quadrant of the tank, and four daily trials were performed for 4 days. On the fifth day, a probe trial was performed similar to that in the acquisition phase. The time spent in the target area and the latency for reaching the target zone were manually measured.

### Recognition memory tasks

The test was adapted from (30). Two visual cues were placed on two adjacent walls of an opaque white Plexiglas cage (58 cm × 50 cm × 43 cm) or in a standard open field arena (UgoBasile, 44 × 44 cm) with black PVC walls and was dimly lit from above (27 lx) the cage. A black and white striped pattern (21 × 19.5 cm) was affixed to the center of the northern wall, and a black and grey checked pattern (26.5 × 20 cm) was placed at the center of the western wall. The stimuli were objects constructed from Duplo blocks (Lego), which varied in shape, color, and size and were too heavy to be displaced. The positions of the objects were randomly counterbalanced between the various positions. All the tasks involved an acquisition phase (5 min) and a recognition test (5 min), separated by a 2- or 24-h delay. A digital camera (Imaging Source, DMK 22AUC03 monochrome) was placed above the apparatus to record the test using a behavioral tracking system (Anymaze 6.0, Stoelting). These videos were used offline by the experimenters blind to the manipulations for *a posteriori* scoring of the time spent in the different zones of the apparatus and exploratory behavior, which was defined as the animal directing its nose toward an object at a distance of at least 2 cm. Climbing or sitting on objects was not considered object exploration. To express the discrimination between the objects, we calculated the discrimination ratio as the absolute difference between the time spent exploring novel, displaced, or exchanged objects and the familiar objects divided by the total time spent exploring the objects.

### Novel object recognitio

In the acquisition phase, two identical objects were placed near the corners (10 cm from the walls). The mice were placed into the arena and allowed to explore for 10 min. One of the two objects was replaced with a novel object during the test. The positions of the objects in the test and those used as novel or familiar were counterbalanced between the mice. *Object location task*. In this task, we measured the ability of mice to recognize an object that had changed location compared to the acquisition phase. In the acquisition phase, two identical objects were placed near the corners. During the test, one object was left in the same position. The other one was displaced to the corner adjacent to the original position, such that the two objects were diagonal from each other. Mice were tested for five minutes.

### Automated visual cue response discrimination and reversal task

The experimental setup was adapted from (6). Experiments were conducted in a standard operant chamber (length of 24 cm × width of 20 cm × height of 18.5 cm; ENV-307W-CT, Med Associates). The chamber was equipped with two nose-poke holes with recessed stimulus lights and a food magazine between them to deliver food rewards (14 mg; test diet, 5-TUL). The setup was placed inside a sound-attenuating cubicle (ENV-022V, Med Associates) homogeneously and dimly lit (6 ± 1 lux) to minimize gradients in light, temperature, sound, and other environmental conditions that could produce a side preference. All tasks were controlled by custom scripts written in MED-PC V (Med Associates). In the SD, the mice were trained to poke into either the nose-poke hole illuminated with green or red lights. Correct responses were reinforced with one food reward, and incorrect responses were signaled with the house light turned on for 5 s. In the simple discrimination reversal (SDRe), reward contingencies were reversed compared to the SD. Animals that learned to respond to the green light had to respond at the red hole to receive a reward and *vice versa*. The mice had to reach a criterion of eight correct choices out of 10 consecutive trials to complete this and each following testing stage. Performance was measured in all phases of all experiments using the number of trials to reach the criterion, time (min) to reach the criterion, and time (s) from breaking the photobeams adjacent to the automated door to a nose-poke response (latency to respond). In addition to this analysis, we classified errors during the SDRe into two subtypes, as done in a previous study (31), to determine whether they reflected problems in shifting from the previously learned strategy (perseverant errors) or maintaining the new strategy after perseveration had ceased (regressive errors). We scored a perseverant error when mice made the same response required on the previous SD on three or more trials per block of four trials. This distinction was made by counting the number of errors made by each mouse in a running window of 12 trials, beginning with trials 1–12 and advancing the window by one trial at a time. When mice made less than three perseverant errors in a block for the first time, all subsequent errors were no longer counted as perseverative errors because, at this point, the mouse chose an alternative strategy at least half of the time. These errors were then scored as regressive errors.

### Fear conditioning

This task was performed as in a previous study (32). Fear conditioning took place in a standard conditioning chamber (Ugo Basile). The conditioned stimulus (CS) was a tone (4 kHz, 80 dB sound pressure level, 30 s), and the unconditioned stimulus (US) was a scrambled shock (0.7 mA; 2 s) delivered through the grid floor that terminated simultaneously with the tone. On experimental day 1, mice were placed in the training chamber. After a 2-min habituation period (baseline), three conditioning trials were presented (tones paired with shock) with an intertrial interval of 90 s. Then, the mice were returned to their home cages 2 min after the last CS–US pairing. After 14 days, mice were re-tested for long-term contextual memory in the same chamber for 5 min without tone or <u>footshock</u>. One hour following the contextual memory, mice were placed in a modified context and, after 2 min of habituation, were exposed to the conditioning tone for 2 min to test for cued memory. Then, the mice were returned to their home cages after 2 min. After another 14 days, mice were re-tested in the conditioned context for 5 min without tone or footshock.

### Neuroanatomical studies

*Systematic neuroanatomical phenotyping* - Morphological studies of the brain were carried out using six homozygous Rnf10-/- (three males and three females, allele Rnf10<tm1b(KOMP)Wtsi) and 12 littermate WT mice (nine males and three females) on a C57BL/6N pure genetic background at 16-weeks of age as previously described (33). Paraffin-embedded brain samples were cut to a 5-μm thickness (sliding microtome, Microm HM 450) to obtain coronal brain region at Bregma +0.98 mm and Bregma −1.34 mm according to the Allen Mouse Brain Atlas (34). Sections were stained with 0.1% Luxol fast blue (Solvent Blue 38; Sigma-Aldrich) and 0.1% cresyl violet acetate (Sigma-Aldrich) and scanned (Nanozoomer 2.0HT, C9600 series) at 20× resolution. *Image analysis* - Sixty-three brain parameters comprised of area and length measurements across the two coronal sections totaling 24 unique brain regions (listed in **Supplementary Table 1**) were used blindly for the genotype. Using in-house ImageJ plugins, an image analysis pipeline was used to standardize measurements of areas and lengths. Images were quality-controlled for the accuracy of sectioning relative to the reference atlas and controlled for asymmetries and histological artifacts. All samples were also systematically assessed for cellular ectopia (misplaced neurons). At Bregma +0.98 mm, the brain structures assessed were as follows: 1. the total brain area, 2. the lateral ventricles, 3. the cingulate cortex, 4. the genu of the corpus callosum, 5. the caudate putamen, 6. the anterior commissure, 7. the piriform cortex, 8. the primary motor cortex, and 9. the secondary somatosensory cortex. At Bregma −1.34 mm, the brain structures were assessed as follows: 10. the total brain area, 11. the lateral and dorsal third ventricles, 12. the retrosplenial granular cortex, 13. the soma of the corpus callosum, 14. the dorsal hippocampal commissure, 15. the dorsal hippocampal region, 16. the amygdala, 17. the piriform cortex, 18. the primary motor cortex, 19. the secondary somatosensory cortex, 20. the mammillothalamic tract, 21. the internal capsule, 22. the optic tract, 23. the fimbria of the hippocampus, and 24. the habenular nucleus. Mice were separately analyzed according to their sex. The data were analyzed using a two-tailed *t*-test assuming equal variance to determine whether a brain region was associated with a neuroanatomical defect. *Neuroanatomical phenotyping of the dorsal hippocampus* - Finer-scale neuroanatomical phenotyping protocols were developed to analyze the hippocampal formation. Additional measurements included the individual CA1, CA2, and CA3 layers and the upper and lower arms of the granular layer of the dentate gyrus. A list of second-level histological parameters is provided in **Supplementary Table 2**.

### DiI labeling for spine morphology

For confocal imaging of the dendritic spines, we labeled neurons with DiI dye (Invitrogen), a fluorescent lipophilic carbocyanine dye, as it diffuses along the neuronal membrane and labels finely dendritic arborization and spine structures in hippocampal slices prefixed with 1.5% PFA. The DiI labeling procedure was performed as previously described (35, 36). In short, DiI solid crystals were applied using a thin needle by lightly touching or gently poking the region of interest on both sides of the 3-mm hippocampal piece, which was prepared after cardiac perfusion with 1.5% PFA in PB 0.1 M. Dil dye was left to diffuse for 1 day in the dark at room temperature (RT). Then, slices were post-fixed with 4% PFA in PB 0.1 M for 45 min at 4°C. The first slice containing the DiI crystals was discarded, and 100-μm hippocampal slices were obtained using a vibratome and collected in PBS. Slices were then mounted on Superfrost glass slides (Thermo Fisher) with Fluoroshield (Sigma) for confocal imaging. Fluorescence images were acquired using the Zeiss Confocal LSM900 system or Nikon A1 Ti2 system with a sequential acquisition setting at 1024 × 1024 pixel resolution. Between 40 and 100 sections of 0.5 μm were acquired for each image, and an appropriate z-projection was obtained.

### Sholl analysis for neuronal branching and dendritic spine morphology

For three-dimensional morphological Sholl analysis, total dendritic length and dendrite morphology were calculated using Fiji freeware software with the Simple Neurite Tracer plug-in. Briefly, a z-stack acquisition was imported, calibrated in Fiji, and semi-automatically traced. The total dendritic length was then computed. The shell interval was set to 5 μm. All analyses were performed blind. In all the experiments, a minimum of five neurons from three independent preparations was analyzed for each condition.

For spine morphology analysis, Z-stack images were analyzed using Fiji (ImageJ) software. Specifically, for each dendritic spine, length, head, and neck width were manually measured at selected regions of interest and then used to classify dendritic spines into three categories (thin, stubby, and mushroom) (37, 38). In particular, the length and ratio between the width of the head and the width of the neck (Wh/Wn) were used as parameters for the classification as follows. Protrusions having a length of more than 3 µm were considered filopodia, and the others were considered spines. Spines with a Wh/Wn ratio above 1.7 were considered mushrooms. Spines with a Wh/Wn ratio smaller than 1.7 were divided into stubby if shorter than 1 µm and thin if longer than 1 µm. Protrusions over 3 µm were qualified as filopodia. Protrusions with lengths over 5 µm were excluded from the analysis. For each neuron, an average of three basal dendrites was considered for a total dendritic length of approximately 200–300 µm. Basal dendrites were analyzed at a distance from the cell body no longer than 300 µm. For *ex-vivo* spine morphology studies, brain samples were analyzed only from mice that had not undergone behavioral and cognitive tests.

### Laser microdissection

*Sample preparation -* PBS-perfused mouse brains were flash-frozen in dry ice-precooled 2-pentane and then cryosectioned into 20-mm-thick slices. The cryosections were collected in pre-cooled MembraneSlide 1.0 PET (Zeiss, Germany) and allowed to dry in the cryostat for 2–3 min at approximately −20°C. For RNA extraction, the cryosections were subsequently incubated in ice-cold 70% ethanol for 2–3 min to reduce RNase activity by dehydration. Before laser microdissection, the sections were left to air-dry for 1–2 min at RT. *Laser capture microdissection -* CA1 and the dentate gyrus were microdissected from the cryosections using a PALM MicroBeam laser microdissector (Zeiss, Germany). Unstained sections were cut using a high laser power energy (20%), and tissue microdissections were collected in *AdhesiveCap* microtubes (Zeiss, Germany). The collected samples were processed immediately for downstream applications (see the Biochemistry and Library preparation sections below).

### Biochemistry

To purify the Triton-insoluble postsynaptic fraction (TIF) highly enriched in postsynaptic density proteins (39), microdissected tissues were homogenized at 4°C in ice-cold buffer (pH 7.4) containing 0.32 M sucrose, 1 mM HEPES, 1 mM MgCl_2_, 1 mM NaHCO_3_ and 0.1 phenylmethanesulfonyl fluoride supplemented with Complete™ protease inhibitor cocktail tablets (Roche Diagnostics) and phosSTOP™ phosphatase inhibitor (Roche Diagnostics). An aliquot of the homogenate was frozen at −20 °C, while the rest of the sample was centrifuged at 1,000 × *g* for 5 min at 4°C to remove nuclear contamination and white matter. The supernatant was collected and spun at 13,000 × *g* for 15 min at 4°C. The resulting pellet (P2-crude membrane fraction) was resuspended in Triton–KCl buffer (1% Triton™ X-100 and 150 mM KCl) and, after 15 min of incubation on ice, spun at 100,000 × *g* for 1 h at 4°C. The pellet (TIF) was resuspended in 20 mM HEPES buffer supplemented with Complete™ protease inhibitor cocktail tablets and stored at −80°C. TIF and homogenate samples for immunoblotting analysis were denatured with Laemmli buffer and 10 min of heating at 98°C. For western blotting assays, the protein contents of TIF and homogenate samples were quantified by Bradford assay. All samples were standardized at a concentration of 1 mg/mL. TIF and total homogenate proteins were separated with SDS–PAGE, followed by western blotting analysis. A total of 10−15 μg of proteins was separated on 6–12% acrylamide/bisacrylamide gel and transferred to a nitrocellulose membrane (Biorad). The membranes were then incubated for 1 h at RT in blocking solution (I-block, TBS 1X, and 20% Tween-20) on a shaker and then incubated with the specific primary antibody in blocking solution overnight at 4°C. The following day, after three washes with TBS and Tween 20 (TBS and 0.1% Tween20; TBSt), they were incubated with the corresponding horseradish peroxidase (HRP)-conjugated secondary antibody in blocking solution for 1 h at RT. After washing with TBSt, membranes were developed with electrochemiluminescence (ECL) reagents (Biorad). Finally, membranes were scanned with a Chemidoc (Biorad Universal Hood III) using Image Lab software (Bio-Rad). Bands were quantified with computer-assisted imaging (Image Lab, Biorad). Protein levels were expressed as relative optical density (OD) measurements normalized to a housekeeping protein.

### RNA extraction and RT–qPCR

Total RNA was extracted from dCA1 microdissected tissues from shRNA RNF10 and control scramble-injected mice (see above for details) with the NucleoSpin, RNA kit (Machery–Nagel) according to the supplier’s specifications. RT–qPCR was performed using the iTaq Universal SYBR Green One-Step Kit (Bio-Rad, 1725151) using the CFX-384 well plate instrument (Bio-Rad). *RNF10* mRNA levels were normalized on *Tubulin a1a* and are expressed as 2^−ΔCt^. The following primers have been used: *RNF10* Forward 5′-CTCACTTTGCTGACCCTGAC-3′, *RNF10* Reverse 5′-CAAGATGGCACTTCATGGCT-3′, *Tubulina1a* Forward 5′-CCGCGAAGCAGCAACCAT-3′, *Tubulina1a* Reverse 5′-CATTGCCGATCTGGACACCA-3′.

### Library preparation

As described above, total RNA was extracted from dCA1 microdissected tissues from shRNA RNF10 and control scramble-injected mice. Libraries were prepared by TruSeq stranded total RNA with Ribo-Zero, and sequencing was performed using an Illumina NextSeq550 platform (paired-end 2 × 75 cycles, ∼40.000.000 reads/sample) in paired-end mode.

### RNA-Seq data analysis

The fastq files generated from the Illumina NextSeq550 were first used to run Multiqc v1.10.1 to assess the overall good sequencing quality. Reads were then mapped to the mouse reference genome GRCm39 with STAR 2.7.9a (40), and Feature Counts 2.0.1 (41) was used to retrieve the counts for each sample. Count data were transformed and normalized for principal component analysis (PCA) to evaluate the sample distribution according to their experimental condition. Differential gene expression analysis was conducted using the Bioconductor package DeSeq2 (42), and genes were considered differentially expressed when they had AdjPval < 0.05 and log2FoldChange ≥ 0.58, indicating a fold change > 1.5 in either direction.

### Ex vivo electrophysiological recordings

*Slice preparation* – Ketamine (100 mg/kg) and xylazine (10 mg/kg) were diluted in saline and injected intraperitoneally in mice 5 min before transcardial perfusion with iced artificial cerebrospinal fluid (aCSF) mixed with a blender. The aCSF contained (all in mM) 120 NaCl, 26 NaHCO_3_, 2.5 KCl, 1.25 NaH_2_PO_4_, CaCl_2_, 1 MgCl_2_, 16.5 glucose, 2.8 pyruvic acid, and 0.5 ascorbic acid, adjusted to pH 7.4 by saturating with carbogen (95% O_2_ and 5% CO_2_), resulting in an osmolarity of 305 mOsm. Parasagittal slices (300 µm) were cut from brain hemispheres using a vibratome (model VT1200S, Leica Microsystems). Slices were then transferred to aCSF for 20 min at 33°C and then maintained at RT until further use.

The recording chamber of the electrophysiology setup was perfused with oxygenated aCSF. Whole-cell patch-clamp recordings were performed from CA1 pyramidal neurons identified with a differential interference contrast microscope (Eclipse FN-1, Nikon, Champigny sur Marne, France) equipped with a camera (CoolSNAP EZ, Ropper, Evry, France). Recordings were performed using borosilicate pipettes (Harvard apparatus: 1.5 OD – 0.86 ID) pulled with a micropipette puller (P97, Sutter Instruments, Novato, CA), with resistances between 4 and 5 MΩ.

Whole-cell voltage-clamp recordings were performed with an internal solution containing (in mM) 125 CsCH_3_SO_3_, 2 MgCl_2_, 4 NaCl, 5 phospho-creatine, 4 Na_2_ATP, 10 EGTA, and 10 HEPES at pH 7.3 adjusted with CsOH and an osmolarity of 300 mOsm. Bicuculline (10 µM) was added to the bath to inhibit GABA-A receptors. Synaptic responses were evoked in CA1 by electrical stimulation (200 µs) of Schaffer collaterals, with electrodes positioned halfway between the hippocampal fissure and the CA1 pyramidal layer with increasing stimulus intensity (20–180 µA). AMPAR and NMDAR EPSCs were recorded at negative (−70 mV) and positive holding potentials (+40 mV), respectively. The NMDAR/AMPAR ratio was calculated as the peak amplitude of the average AMPAR EPSCs (10 consecutive events) divided by the average peak amplitude of NMDAR EPSCs (10 consecutive events).

Whole-cell current-clamp recordings were performed with an internal solution containing (in mM): 140 K-methanesulphonate, 2 MgCl_2_, 10 HEPES, 0.3 EGTA, 2 MgATP, and 0.3 NaGTP at pH 7.3 adjusted with KOH and an osmolarity of 300 mOsm. To investigate the intrinsic membrane and firing properties of neurons, we applied 500-ms depolarizing current steps from −80 pA to +320 pA in increments of 20 pA. The hyperpolarizing step was used to calculate the sag, the membrane capacitance (Cm), the input resistance (Rin), and the time constant.

fEPSP were recorded in the stratum radiatum of CA1. Tetanic-burst stimulation (five bursts of four stimuli at 100 Hz separated by 200 ms) was triggered to induce long-term potentiation. To generate summary graphs, individual experiments were normalized to the baseline and averaged to generate 30-s bins. These were then averaged to generate the final summary graphs. The magnitude of LTP was calculated based on the fEPSP values 50-60 min after the end of the induction protocol.

Signals were amplified using a HEKA EPC10 amplifier (Lambrecht, Germany), filtered at 3.3 kHz, and digitized at 10 kHz via PatchMaster software (Lambrecht, Germany). Data were analyzed using IGOR PRO 6.3 (Wavemetrics, Lake Oswego, OR) and Neuromatic version 2.6 software.

### Antibodies and other reagents

Rabbit anti-GluA1 (13185S, Cell Signaling Technology, dilution: 1 : 1,000 WB); rabbit anti-pGluA1^S845^ (3420, Abcam, dilution: 1 : 1,000 WB); rabbit anti-pERK (9101, Cell Signaling Technology, 1 : 1000), rabbit anti-ERK (9102, Cell Signaling Technology, 1 : 1000), rabbit anti-RASGRF2 (PA5-28867, Invitrogen, 1 : 1000), rabbit anti-NMDAR2A (M264, Sigma, 1 : 1000), rabbit anti-pCREB (9198, Cell Signaling Technology, 1 : 1000), rabbit anti-CREB (9197, Cell Signaling Technology, 1 : 1000); mouse anti-Tubulin (T9026, Sigma-Aldrich, dilution: 1 : 5,000 WB). The following secondary antibodies were used: goat anti-mouse-HRP (172–1011, Bio-Rad, dilution: 1 : 10,000); goat anti-rabbit-HRP (170–6515, Bio-Rad, dilution: 1 : 10,000); goat anti-rabbit-Alexa488 (A-11034, Invitrogen, 1 : 1000); and goat anti-mouse-Alexa555 (A-21424, Invitrogen, dilution: 1 : 1000).

### Quantification and statistical analysis

Statistical analysis was performed with GraphPad Prism software, and data are presented as means ± s.e.m. (standard error of the mean). For molecular, morphological, and behavioral analysis, normal distribution was checked using the D’Agostino & Pearson normality test or Shapiro–Wilk normality test. The tests used to assess data significance are indicated in the figure legends. We used a two-tailed unpaired *t*-test (a p-value less than 0.05 was considered significant), one- or two-way ANOVA as appropriate, followed by Tukey or Bonferroni’s test as a post-hoc test. Images acquired with a confocal microscope were analyzed using Fiji/Image J software.

## RESULTS

### RNF10 loss impairs hippocampal plasticity and cognitive flexibility

To determine whether RNF10 loss alters functional synaptic responses, we examined the electrophysiological properties of hippocampal CA1 neurons from RNF10 KO mice. Whole-cell recordings (**Figure 1A-B**) revealed increased intrinsic excitability in RNF10-deficient neurons, characterized by a significant reduction in rheobase (**Figure 1C**) and firing threshold (**Figure 1D**), together with an elevated firing rate compared with wild-type (WT) littermates (**Figure 1E**), while sag amplitude remained unchanged (**Figure 1F-G**). Consistent with these alterations, field recordings in the CA1 stratum radiatum (**Figure 1K**) showed that tetanic-burst stimulation induced robust long-term potentiation (LTP) in WT mice but failed to do so in RNF10 KO animals, indicating a marked impairment of activity-dependent synaptic plasticity (**Figure 1L-O**). In contrast, basal synaptic transmission was preserved: the frequency and amplitude of spontaneous synaptic events were unchanged (**Supplementary Figure 1A–E**), stimulation of Schaffer collaterals produced similar AMPAR- and NMDAR-mediated responses (**Supplementary Figure 1F–H**), and the NMDAR/AMPAR ratio was not altered between genotypes (**Supplementary Figure 2I**). Passive membrane properties, including membrane capacitance (**Figure 1H**), input resistance (**Figure 1I**), and membrane time constant (**Figure 1J**), were also comparable between RNF10 KO and WT neurons. Together, these findings indicate that RNF10 loss selectively disrupts intrinsic excitability and long-term synaptic plasticity in CA1 neurons while sparing basal synaptic transmission.

**Figure 1.**
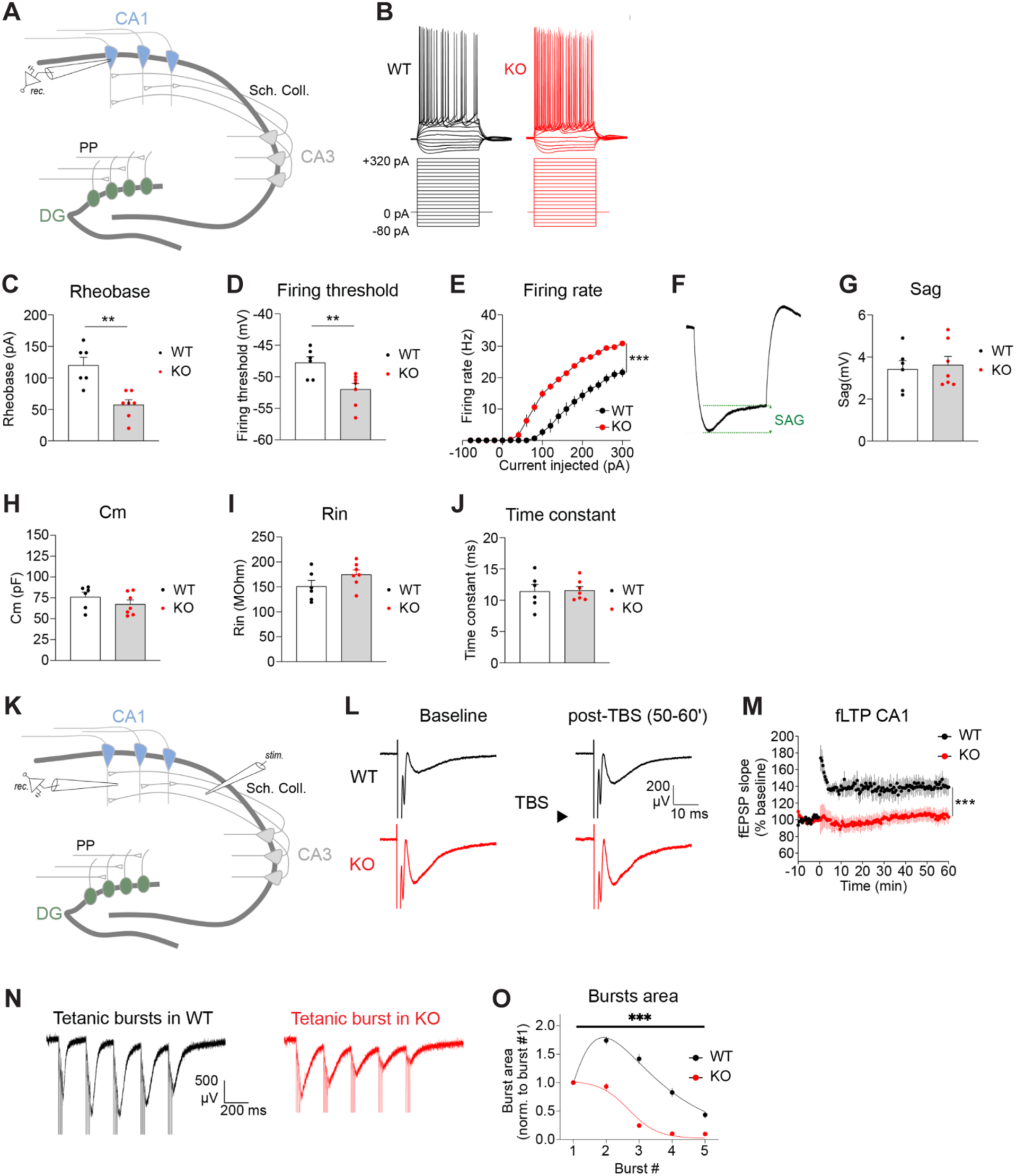
Increased excitability and impaired long-term plasticity in CA1 neurons of RNF10 KO mice. (**A**) Schematic of the main hippocampal sub-regions. (**B-J**) Results were obtained by recording the intrinsic membrane and firing properties of CA1 pyramidal neurons. (**B**) Electrophysiological recording representing the membrane potential during injection of negative or positive current steps for 500 ms. (**C, D**) The rheobase and firing threshold significantly decreased with the absence of RNF10 (H; two-tailed unpaired *t*-test, p = 0.0012. I; two-tailed unpaired *t-*test, p = 0.0099). (**E**) The firing rate significantly increased with absence of RNF10 (Two-way ANOVA, Current injected, F(_19, 220_) = 171.1, p < 0.0001; Genotype, F(_1, 220_) = 391.0, p < 0.0001, Interaction, F (19, 220) = 10.98, p < 0 .0001). (**F**) Electrophysiological trace representing the sag induced by hyperpolarizing CA1 neurons. (**G**) The sag amplitude is similar between RNF10 WT and KO (two-tailed unpaired *t*-test, p = 0.7418). (**H–J**) The membrane capacitance (Cm), resistance (Rin), and the time constant of CA1 neurons were similar between WT and RNF10 KO mice. (E; two-tailed unpaired *t*-test, p = 0.2691. F; two-tailed unpaired *t*-test, p = 0.1376. G; two-tailed unpaired *t*-test, p = 0.8940). (**K**) Schematic of the main hippocampal sub-regions. (**L–O**) Results were obtained by recording field excitatory post-synaptic potentials (fEPSP) in the CA1 stratum radiatum. (**L**) Electrophysiological recordings representing the fEPSP before (baseline) and after (post-TBS) tetanic burst stimulation (TBS). The fEPSP was potentiated in WT but not in RNF10 KO mice. (**M**) Analysis of the fEPSP slope before and after TBS triggered at time 0. The fEPSP was potentiated in WT but not in RNF10 KO mice (two-tailed unpaired *t*-test, WT baseline vs. 50–60 min, p = 0.0010; RNF10 KO baseline vs. 50–60 min, p = 0.5981). (**N**) Electrophysiological recording representing the extracellular potential during bursts of high-frequency stimulations (five bursts of four stimuli at 100 Hz separated by 200 ms). (**O**) Analysis of the priming effects of repetitive burst stimulation. The area under bursts #2 and #3 increased in WT, whereas the area under the bursts decreased over repetitive stimulation in the KO mice (Two-way ANOVA, burst number, F(_4, 630_) = 126.0, p < 0.0001; genotype, F (1, 630) = 322.6, p < 0.0001; interaction, F(_4, 630_) = 35.5, p < 0.0001). **p < 0.01, ***p < 0.005. Values are expressed as means ± s.e.m.

We next asked whether alterations in hippocampal plasticity translate into deficits in flexible updating upon contingencies change. We first assessed cognitive flexibility in RNF10 KO mice WT littermates using a Morris water maze paradigm that includes a reversal phase. While the acquisition phase measures learning and memory, the reversal stage requires animals to suppress a previously learned spatial strategy and update their behavior when contingencies change. In the acquisition phase, RNF10 KO mice showed a longer latency to reach the target area than control WT mice only in the late phase of the test (day 3) (**Figure 2A**). However, following the acquisition, RNF10 KO mice displayed normal spatial memory (probe), as indicated by the latency to reach the target area and time spent in this area (**Figure 2B, C**), compared to WT mice. Importantly, when we changed the position of the target area to the opposite quadrant of the arena to test the reversal learning abilities of the mice, RNF10 KO mice displayed a large increase in latency to reach the target area compared to controls throughout the task, suggesting a deficit in reversal learning (**Figure 2D**). In line with these results, RNF10 KO mice also displayed a reduced time spent in the target area during the probe test, following reversal learning, suggesting they could not properly recall the location of the target area (**Figure 2E, F**). These results suggest that RNF10 deficiency may impair cognitive flexibility.

**Figure 2.**
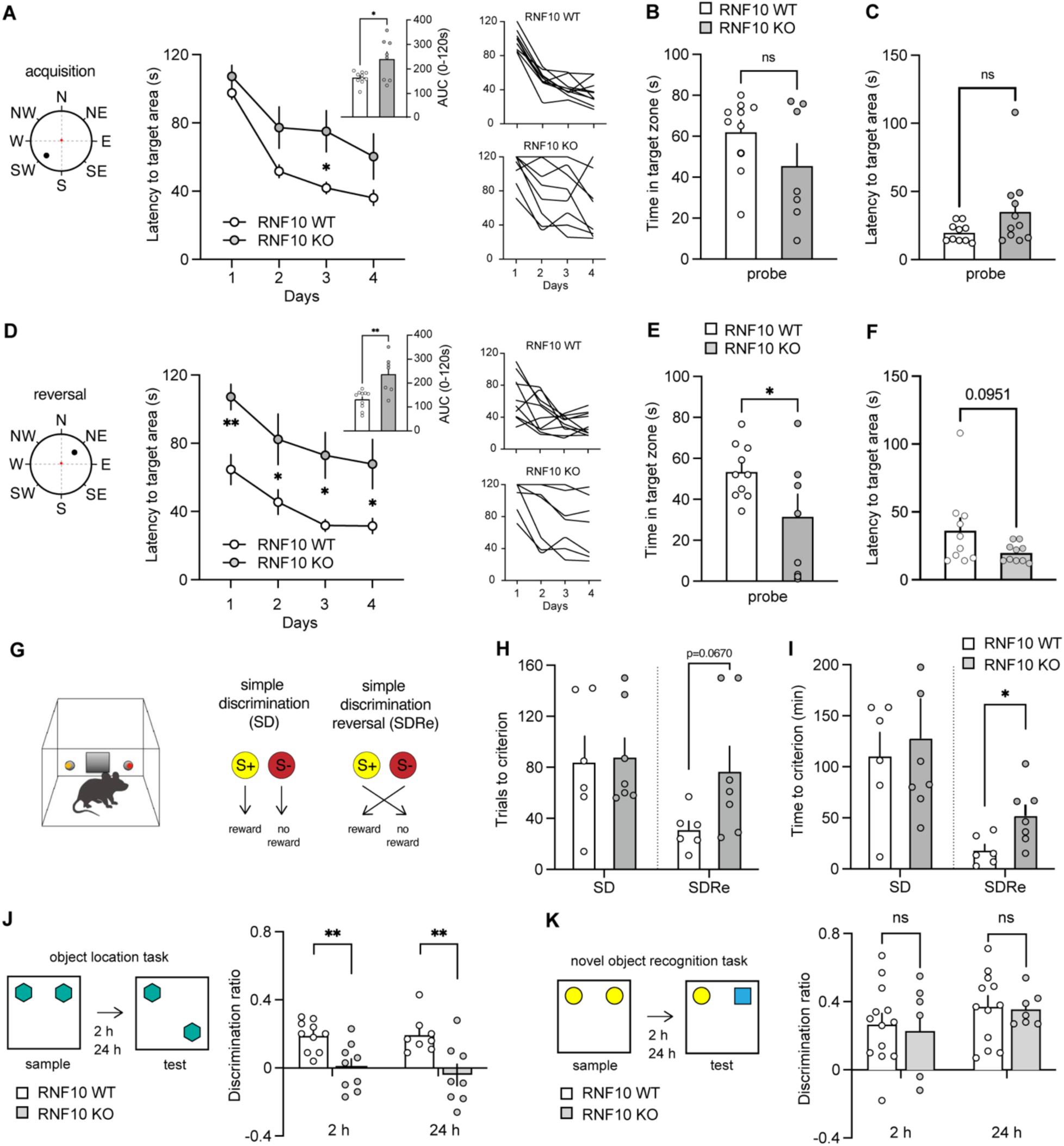
RNF10 KO impairs spatial and reversal learning. (**A**) *Left*, latency to reach the target area during the 4 days of acquisition of the Morris water maze (two-way RM ANOVA, time, F(_3,48_) = 39.38, p < 0.0001; genotype, F(_1,16_) = 7.08, p = 0.0171) in WT (n = 10) and RNF10 KO (n = 8) mice (Inset, area under the curve (AUC) in the latency to target area, two-tailed unpaired *t*-test, t = 2.71, df = 16, p = 0.0153). *Right*, individual curves of latency to target area. (**B**) Time in the target zone (two-tailed unpaired *t*-test, p = 0.1627) and (**C**) latency to target area (two-tailed unpaired *t*-test, p = 0.1627) during the probe test 24 h following the last trial in the Morris water maze in WT (n = 10) and RNF10 KO (n = 11) mice. (**D**) *Left*, latency to reach the target area during the 4 days of the reversal stage of the Morris water maze (two-way RM ANOVA, time, F(_3,45_) = 15.84, p < 0.0001; genotype, F(_1,15_) = 12.97, p = 0.0026) in WT (n = 10) and RNF10 KO (n = 8) mice (Inset, AUC in the latency to target area, two-tailed unpaired *t*-test, p = 0.0153). *Right*, individual curves of latency to target area. (**E**) Time in the target zone (two-tailed unpaired *t*-test, p = 0.1627) and (**F**) latency to target area (two-tailed unpaired *t*-test, p = 0.1627) during the probe test 24 h following the last trial in the reversal phase of the Morris water maze in WT (n = 10) and RNF10 KO (n = 10) mice. (**G**) Schematic of the automated visual cue simple discrimination (SD) and reversal task. In the SD, mice were presented with a two-choice decision-making task, where a visual cue, randomly assigned to a left or right nose poke, was associated with a food reward (S+). The other stimulus did not result in reward delivery and was associated with house light on (S−). In the simple discrimination reversal (SDRe), mice were required to reverse their response. Mice had to choose the previously irrelevant visual cue to receive a reward. (**H**) Number of trials (SD: two-tailed unpaired *t*-test, p = 0.8745; SDRe: two-tailed unpaired *t*-test, p = 0.0670) and **(I)** time (SD: two-tailed unpaired *t*-test, p = 0.7226; SDRe: two-tailed unpaired *t*-test, p = 0.0263) required by the WT (n = 6) and RNF10 KO (n = 7) mice to complete the SD and SDRe. (**J**) *Left*, Schematic of the object location test. *Right*, performance in the object location test 2 h and 24 h following the sample phase of WT (n = 10/8) and RNF10 KO (n = 9/8) mice, expressed as the discrimination ratio (two-tailed unpaired *t*-test, 2 h: p = 0.0058, 24 h: p = 0.0065). (**K**) *Left*, Schematic of the novel object recognition test. *Right*, performance in the novel object recognition task 2 h and 24 h following the sample phase for WT (n = 13/12) and RNF10 KO (n = 7) mice, expressed as the discrimination ratio (two-tailed unpaired *t*-test, 2 h: p = 0.9315, 24 h: p = 0.9315). All values are expressed as means ± s.e.m. *p < 0.05, **p < 0.01, ns, not significant.

To understand whether reversal learning deficit of RNF10 KO mice was specific for navigational features or could be extended to different non-spatial cues, we designed an instrumental reversal learning task to assess flexibility of goal-directed behavior (**Figure 2G)**. In this task, mice were presented with two nose-poke holes equipped with recessed light cues switched on in pairs and counterbalanced between left and right nose-poke holes. Mice were first trained in a two-choice simple discrimination (SD) to poke into either left or right nose-poke holes to receive a reward pellet. Thus, the discriminative association between the correct response, resulting in food delivery, and the nose-poke hole varied depending on a visual cue. Mice reached a criterion of eight correct choices out of 10 consecutive trials to complete the SD. Then, they were tested in the reversal learning stage (SDRe), where the reward contingencies were reversed. Both control and RNF10 KO mice readily acquired the SD. However, RNF10 KO mice required more trials (**Figure 2H)** and more time (**Figure 2I)** to solve the reversal stage than their control littermates. Altogether, these results suggest that RNF10 KO mice might have an impaired ability to disengage from acquired behavior and display an adaptive behavioral response following new contingencies across spatial and non-spatial domains.

We next examined performance in the object location task, which requires updating of spatial contextual representations, implying major hippocampus involvement (43). In this task, after familiarization with two identical objects, one of the two was displaced to the opposite corner of the arena (**Figure 2J**). In this test, RNF10 KO mice displayed lower scores of discrimination ratio 2 and 24 hours following the sample phase (**Figure 2J**). As control, we also tested mice in the novel object recognition task (**Figure 2K**) in which the mice recognize the newly introduced object based on familiarity. In this case, both WT and RNF10 KO mice were able to discriminate the objects 2 and 24 hours following the familiarization phase (sample) (**Figure 2K**). This suggests that deficit in recognition memory was not generalized but specific for spatial features.

### RNF10 signaling in dorsal CA1 prevents perseveration and supports cognitive flexibility

Given the prominent role of hippocampal circuits in context-dependent learning (11, 12) and contingency-driven updating (9), we next asked whether lack of cognitive flexibility observed in RNF10 KO mice could be attributed to dysfunction within dorsal hippocampal networks. Consistent with this hypothesis, neuroanatomical analyses in RNF10 KO mice revealed a marked microcephalic phenotype with a significant reduction in total brain area and prominent hypoplasia of the dorsal hippocampus, accompanied by commissural abnormalities. These alterations were comparable in males and females and were particularly evident at Bregma −1.34 mm, where the dorsal hippocampus and dentate gyrus showed significant size reductions (**Supplementary figure 2A-C)**. Together, these findings provide convergent evidence that RNF10 loss disrupts hippocampal organization. We therefore targeted RNF10 expression in the dorsal CA1 (dCA1) of adult mice using viral shRNA (ShRNF10) to test whether hippocampal RNF10 signaling is sufficient to control behavioral updating, while minimizing potential developmental compensation associated with constitutive gene deletion. As control, we injected a virus carrying a scramble sequence (Scr) (**Figure 3A**).

**Figure 3.**
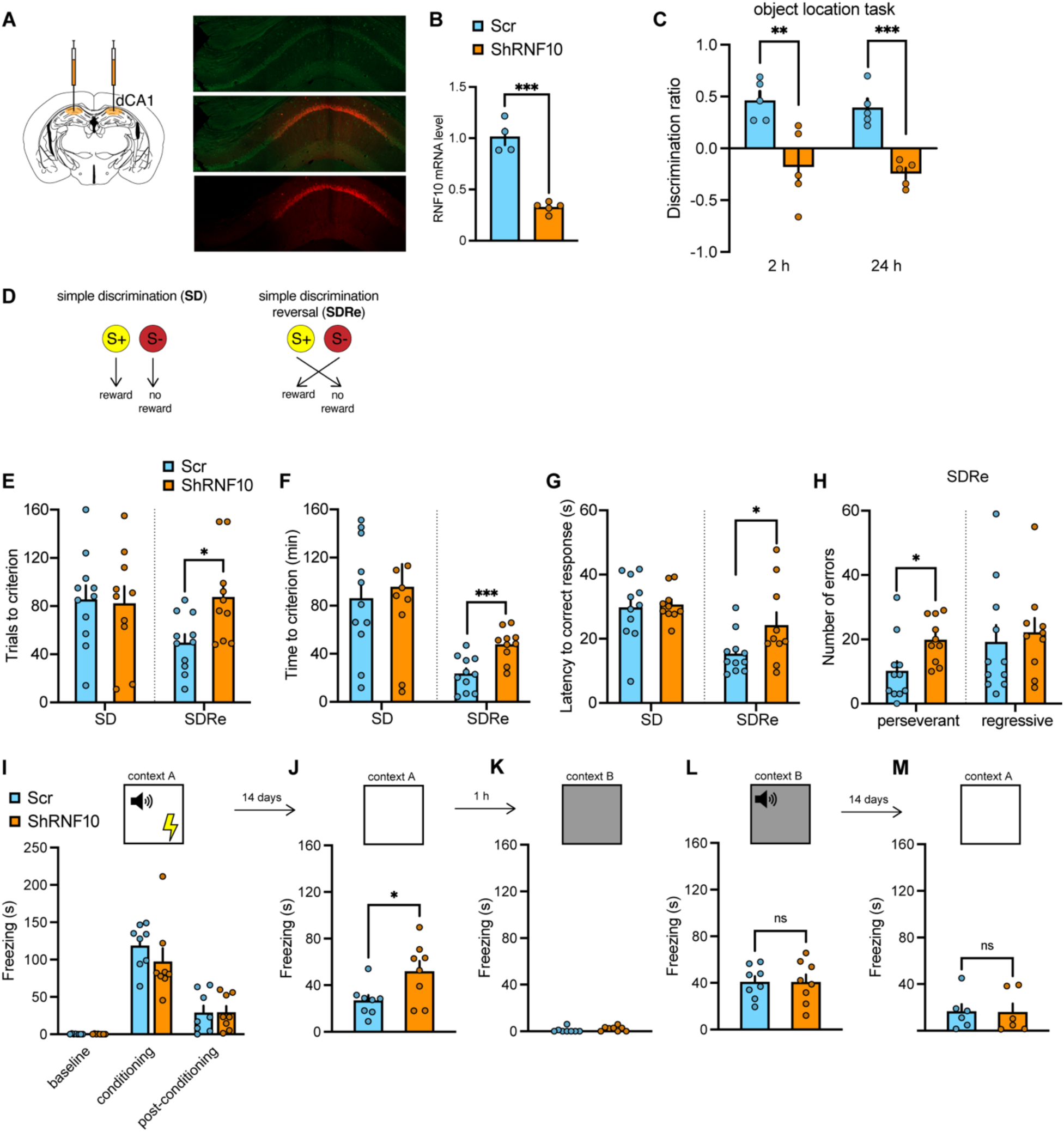
RNF10 silencing in the dCA1 recapitulate deficits of reversal learning. (**A**) Male mice were bilaterally injected in the dCA1 with AAV carrying specific short hairpin RNAs (shRNF10, orange) to silence endogenous RNF10 or scramble shRNA (Scr, light blue). Representative image of a coronal section of dCA1. (**B**) mRNA levels of endogenous RNF10 following injection with shRNA RNF10 or a scramble shRNA in the dCA1 (two-tailed unpaired *t*- test, p < 0.001, n = 4/5). (**C**) Performance in the object location test 2 and 24 h following the sample phase for Scr (n = 5) and ShRNF10 (n = 5) mice, expressed as the discrimination ratio (two-tailed unpaired *t*-test, 2 h: p = 0.0081, 24 h: p = 0.0004). (**D**) Schematic of the automated visual cue response discrimination and reversal test. (**E**) Number of trials (SD: two-tailed unpaired *t*-test, p = 0.8650; SDRe: two-tailed unpaired *t*-test, p = 0.0112), (**F**) Time (SD: two-tailed unpaired *t*-test, p = 0.6923; SDRe: two-tailed unpaired *t*-test, p = 0.0007), and (**G**) Latency to make a correct response (SD: two-tailed unpaired *t*-test, p = 0.8090; SDRe: two-tailed unpaired *t*-test, p = 0.050) required by Scr (n = 11) and ShRNF10 (n = 10) mice to complete the SD and SDRe. (**H**) Number of perseverant (two-tailed unpaired *t*-test, t = 2.53, df = 19, p = 0.020) and regressive (two-tailed unpaired *t*-test, p = 0.6690) errors made by Scr (n = 11) and ShRNF10 (n = 10) mice during the SDRe. (**I**) Freezing behavior (expressed in s) of Scr (n = 8) and ShRNF10 (n = 8) mice during baseline, conditioning (three-tone–shock pairings), and post-conditioning stages of the fear conditioning learning (two-way RM ANOVA, stage x group, F(_2,28_) = 0.73, p = 0.4893). (**J**) Freezing behavior of Scr (n = 8) and ShRNF10 (n = 8) mice during memory recall 2 weeks following the conditioning in the conditioning chamber (context A, two-tailed unpaired *t*-test, p = 0.0268) or in (**K**) a modified chamber (context B, two-tailed unpaired *t*-test, p = 0.2474) or in (**L**) a modified chamber in the presence of the conditioned tone (two-tailed unpaired *t*-test, p = 0.9856). (**M**) Freezing behavior of Scr (n = 6) and ShRNF10 (n = 6) mice during the memory recall 4 weeks following conditioning (two-tailed unpaired *t*-test, p = 0.9587). *p < 0.05, **p < 0.01, ***p < 0.005. Values are expressed as means ± s.e.m.

The shRNA against RNF10 significantly reduced the expression of RNF10 in the dCA1 (**Figure 3B**). Four weeks following viral injection, control mice and those with RNF10 silencing in dCA1 were assessed for their cognitive abilities. We tested the mice in the object location task. We found that mice injected with shRNF10 did not discriminate the displaced object (**Figure 3C**), similar to what was observed in RNF10 KO mice (**Figure 2J**).

Using the operant two-choice SD task, we then investigated whether RNF10 silencing in the dCA1 could affect the reversal learning ability (**Figure 3D**). The RNF10-silenced mice required more trials (**Figure 3E**) and time (**Figure 3F**) than controls to solve the reversal stage (SDRe) but not the initial discrimination task (SD). This effect phenocopies the one found in RNF10 KO mice (**Figure 2H, I)**. In the reversal stage (SDRe), RNF10-silenced mice required more time to respond correctly (**Figure 3G**). We then classified errors to understand whether RNF10-silenced mice had an altered ability to shift from previously learned discrimination (perseverative errors) or maintain a new strategy after perseveration finished (regressive errors). This analysis showed that in the reversal stage, RNF10-silenced mice made more perseverant, but not regressive, errors than controls (**Figure 3H**). Thus, these results suggest that silencing RNF10 in the dCA1 of mice alters their ability to adapt their responses to changing situational demands flexibly

To understand whether the increased cost in flexibility could be related to the stability of the prior acquired information, we tested shRNF10 and control mice in a fear-conditioning paradigm that allows the formation of long-lasting memories following one training session (44). We found that both groups of mice similarly acquired fear learning (**Figure 3I**). However, when mice were re-exposed to the conditioning context 14 days following the acquisition, the RNF10-silenced mice showed more freezing behavior than controls (**Figure 3J**). This behavior was specific to the conditioning context, as freezing behavior disappeared during exposure to a different context (**Figure 3K**). Presentation of the conditioned stimulus elicited freezing behavior, but we did not find any differences between groups (**Figure 3L**). Finally, we tested the mice again in the conditioning context following another 14 days, and we found an expected weakening of freezing behavior in both RNF10-silenced and control mice (**Figure 3M**). Altogether, these results suggest that silencing of RNF10 in the dCA1 strengthens the initial acquisition of learning that becomes detrimental when adaptation of behavior is required when challenged with a changing context.

To determine whether the behavioral alterations observed following RNF10 silencing were attributable to loss of RNF10, we restored RNF10 expression in the dCA1 by injected bilaterally a 1:1 mixture of viruses for the expression of the shRNF10 and an ShResistant-RNF10 (ShRNF10 + ShResistant) . Overall, these mice showed the concomitant expression of both viral vectors that ultimately brought to RNF10 increased expression in the shRNF10+ShResistant group **(Figure 4A).** Interestingly, these mice showed a complete rescue in the ability to discriminate the displaced object in the object location task at 24 hours, further suggesting the role of RNF10 in spatial recognition memory (**Figure 4B**).

**Figure 4.**
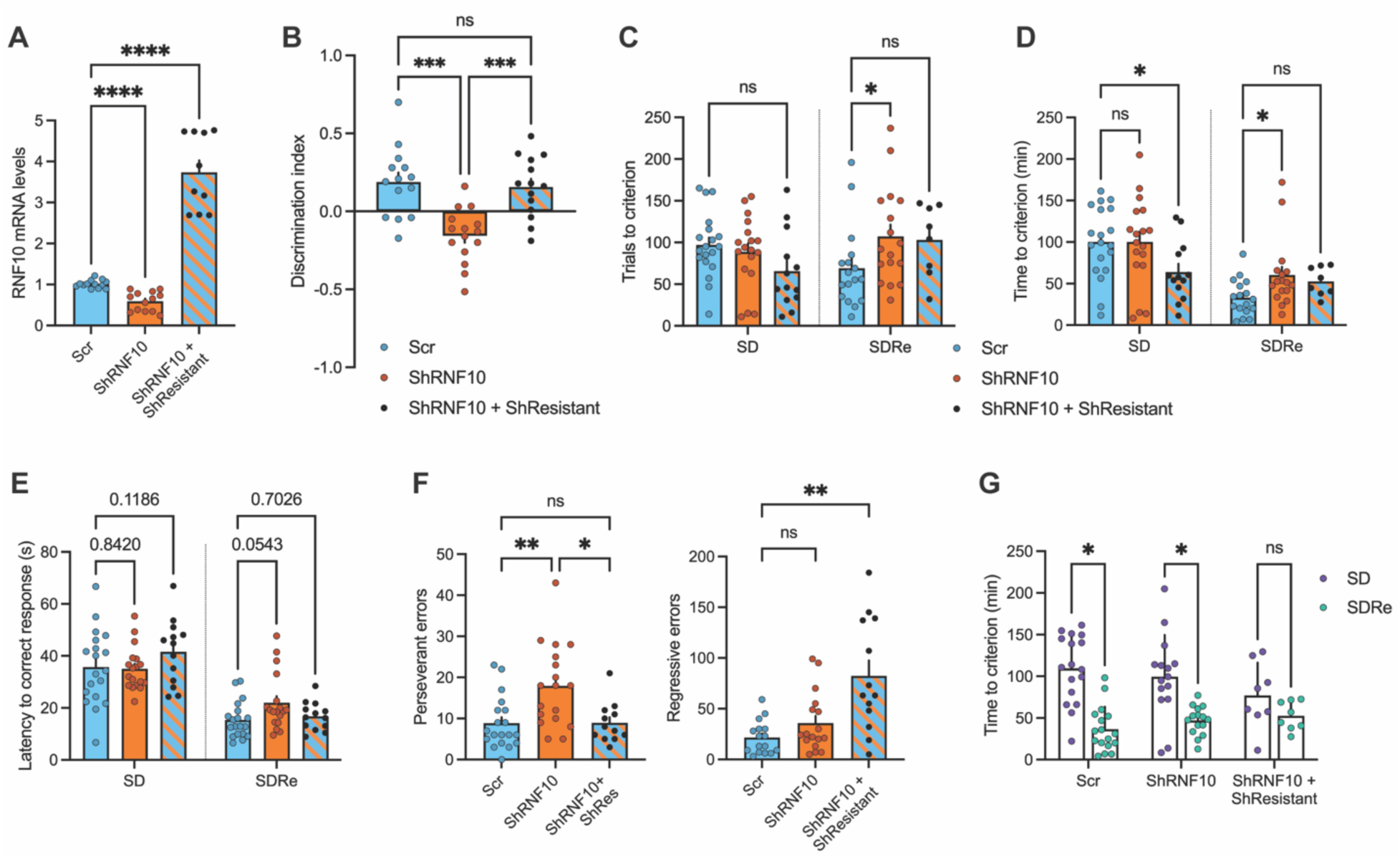
Rescue of cognitive flexibility in ShRNF10+ShResistant mice. (**A**) qRT-PCR analysis for the expression of endogenous RNF10 following injection with scramble shRNA (n=13), shRNF10 (n=13) or shRNF10 + ShResistant (n=10) in the mouse dCA1. Gene expression was normalized to *TUBA1A* (Brown–Forsythe ANOVA with Dunnett’s T3 multiple comparisons test. p≤0.0001). (**B**) Performance in the object location test 24 h following the sample phase for Scr (n = 14), ShRNF10 (n = 15) and ShRNF10 + ShResistant (n = 14) mice, expressed as the discrimination ratio (One-way ANOVA with Tukey’s post hoc test. Scr vs ShRNF10 p=0.0001; Scr vs Sh+ShResistant p=0.9078; ShRNF10 vs Sh+ShResistant p=0.0004) (**C**) Number of trials (Mixed-effects model with Fisher’s LSD multiple comparisons test. SD: Scr vs ShRNF10 p=0.5918; Scr vs Sh+ShResistant p=0.0656; ShRNF10 vs Sh+ShResistant p=0.1781; SDRe: Scr vs ShRNF10 p=0.0202; Scr vs Sh+ShResistant p=0.1125; ShRNF10 vs Sh+ShResistant p=0.7725) and (**D**) time (Mixed-effects model with Fisher’s LSD multiple comparisons test; single pooled variance. SD: Scr vs ShRNF10 p=0.9953; Scr vs Sh+ShResistant p=0.0119; ShRNF10 vs Sh+ShResistant p=0.0130; SDRe: Scr vs ShRNF10 p=0.0456; Scr vs Sh+ShResistant p=0.2449; ShRNF10 vs Sh+ShResistant p=0.6527) required by Scr (n = 19/17), ShRNF10 (n = 18/17) and ShRNF10 + ShResistant (n = 13/8) mice to complete the SD and SDRe. (**E**) Latency to make a correct response (Mixed-effects model with Fisher’s LSD multiple comparisons test; single pooled variance; SD: Scr vs ShRNF10 p=0.8420; Scr vs Sh+ShResistant p=0.1186; ShRNF10 vs Sh+ShResistant p=0.0855; SDRe: Scr vs ShRNF10 p=0.0543; Scr vs Sh+ShResistant p=0.7026; ShRNF10 vs Sh+ShResistant p=0.1701) required by Scr (n = 19/17), ShRNF10 (n = 18/17) and ShRNF10 + ShResistant (n = 13/8) mice to complete the SD and SDRe. (**F**) Number of perseverant and regressive errors made by Scr (n = 19), ShRNF10 (n = 18) and ShRNF10 + ShResistant (n = 13) mice during the SDRe (Kruskal–Wallis test with Dunn’s multiple comparisons test. Perseverant: Scr vs ShRNF10 p=0.0054; Scr vs Sh+ShResistant p>0.9999; ShRNF10 vs Sh+ShResistant p=0.0358; Regressive: Scr vs ShRNF10 p=0.0.6805; Scr vs Sh+ShResistant p=0.0016; ShRNF10 vs Sh+ShResistant p=0.0527). (**G**) Comparison of time required to complete the SD vs SDRe for each group of mice (Wilcoxon test with Holm–Šidák multiple comparisons correction. Scr p<0.0001; ShRNF10 p=0.0201; Sh+ShResistant p=0.1484). Values are expressed as means ± s.e.m. *p < 0.05, **p < 0.01, ***p < 0.001, ****p<0.0001.

We next assessed whether RNF10 re-expression could rescue deficits in reversal learning. ShRNF10+ShResistant mice required significantly less time and trials to reach criterion in the initial SD task, indicating enhanced acquisition of the stimulus–reward association in the first phase (**Figure 4C, D**). During the reversal stage (SDRe), they required a similar number of trials and time as ShRNF10 mice (**Figure 4C, D**). Notably, ShRNF10+ShResistant mice showed a modest tendency toward normalization of the latency to respond correctly in the SDRe (**Figure 4E**). Interestingly, error pattern analyses revealed that ShRNF10 + ShResistant mice displayed levels of perseverative errors comparable to controls, consistent with a rescue of cognitive flexibility. In contrast, they exhibited an increased number of regressive errors compared to both control and ShRNF10 mice (**Figure 4F**). While animals typically showed a reduction in completion time from SD to SDRe, this effect was absent in ShRNF10+ShResistant mice (**Figure 4G**). Together, these results indicate that although overall reversal performance was not fully restored, RNF10 re-expression abolished perseverative responding and induced an alternative behavioral strategy adopted during contingencies updating, suggesting that animals processed the reversal stage as a novel discrimination rather than as a modification of the previously acquired rule. Overall, these findings indicate that restoring RNF10 expression in the dCA1 modifies the behavioral phenotype induced by RNF10 silencing and supports a role for RNF10 signaling in regulating cognitive flexibility and the ability to disengage from previously learned strategies.

### RNF10 depletion leads to structural and functional alterations in the hippocampus

RNF10 silencing has been described in *in vitro* systems to induce structural modification of dendritic spines (26) and dendritic branching (27). Because morphological spines’ dynamic is a key determinant of activity-dependent plasticity, we asked whether similar alterations occur *in vivo* following RNF10 depletion in the hippocampus. We performed DiI-staining of hippocampal slices from shRNF10 and Scr mice four weeks after viral injection to analyze dendritic spines (**Figure 5A**). Dendritic spines of dCA1 neurons from shRNF10 mice showed a significant reduction in both spine width (**Figure 5B**) and length (**Figure 5C**), with no difference in the total number of protrusions (**Figure 5A**). Of notice, similar results were found on hippocampal slices of adult RNF10 KO and WT mice (**Figure 5D**). Indeed, hippocampal CA1 pyramidal neurons in RNF10 KO mice showed a significant reduction in both spine head width (**Figure 5E**) and spine length (**Figure 5F**), with no changes in spine density (**Figure 5D**). These results support a specific role of RNF10 in regulating dendritic spine morphology in CA1 neurons.

**Figure 5.**
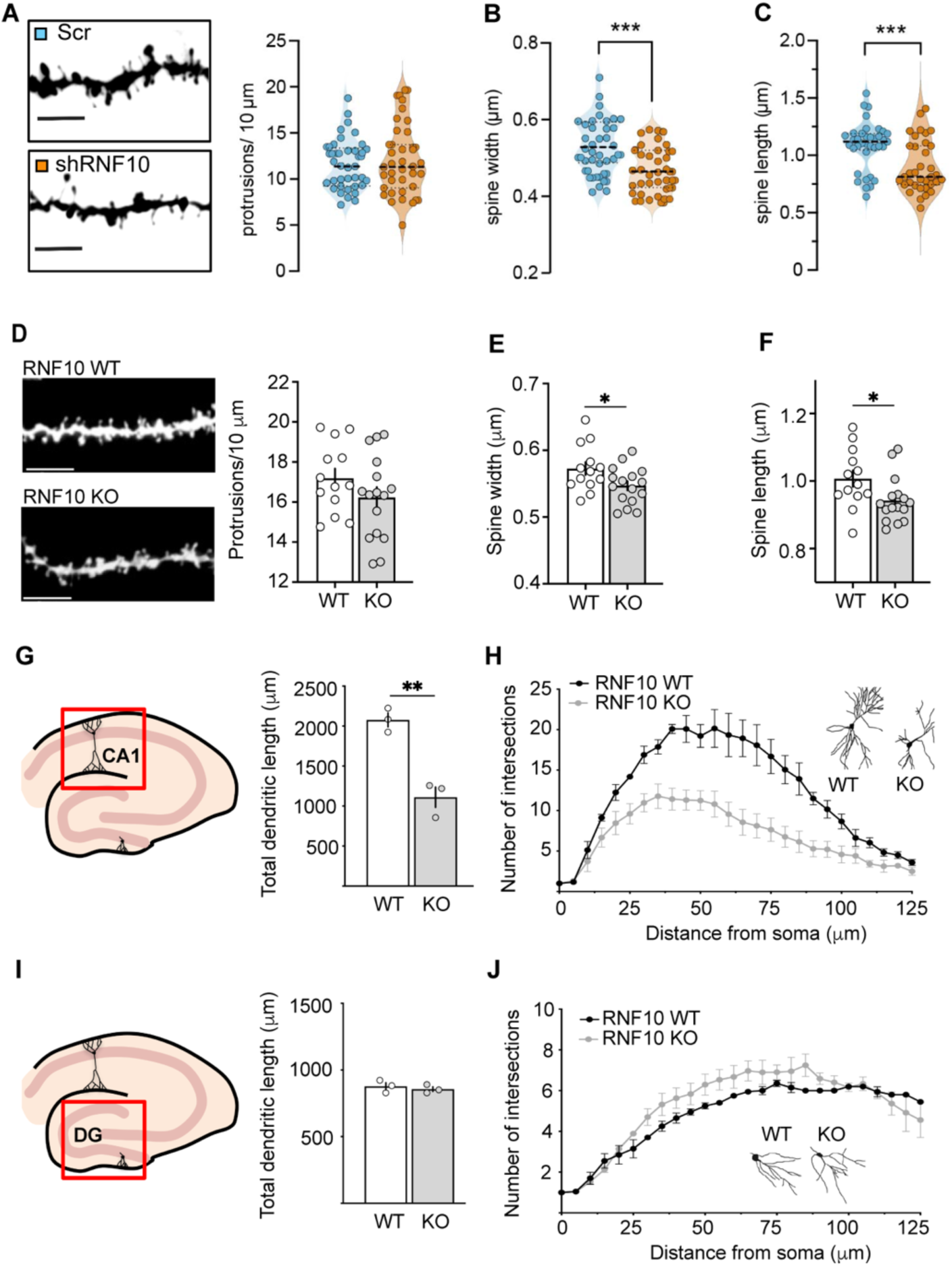
RNF10 loss alters dendritic spine morphology in dCA1. **(A)** Representative images showing dendrites of adult mice dCA1 neurons and protrusion densities (two-tailed unpaired t-test p = 0.7367, n = 37/35) after injecting either RNF10 shRNA or the scramble control (scr; scale bar = 5 μm). (**B**) Violin plots representing, for both conditions dendritic spine width (two-tailed unpaired t-test; p = 0.0002, n = 36/35), and (**C**) dendritic spine length (two-tailed unpaired t-test; p = 0.0006, n = 37,35). (**D**) Representative images showing dendrites of adult mice dCA1 neurons of adult RNF10 KO and WT mice (scale bar = 5 μm) and quantification of protrusion densities (two-tailed unpaired t-test, p = 0.1935, n = 13/16). (**E**) Bar graphs representing, for both conditions, dendritic spine width (two-tailed unpaired t-test, p = 0.0353, n = 13/16) and (**F**) dendritic spine length (two-tailed unpaired t-test, p = 0.0327, n = 13/16). (**G**) Total dendritic length (two-tailed unpaired t-test, p = 0.0033, n = 3) and (**H**) representative sketched neurons and quantification via Sholl analysis (two-tailed paired t-test, p < 0.0001) of CA1 hippocampal neurons from brain slices of adult RNF10 KO and WT mice. (**I**) Total dendritic length (two-tailed unpaired t-test, p = 0.5581, n = 3) and (**J**) representative sketched neurons and quantification via Sholl analysis (F; two-tailed paired t-test, p = 0.5810) of DG hippocampal neurons from brain slices of adult RNF10 KO and WT mice. *p < 0.01, **p < 0.005, ***p<0.001. Values are expressed as means ± s.e.m.

To further examine whether RNF10 deficiency alters dendritic architecture, we next performed Golgi staining on hippocampal slices of adult RNF10 KO and WT mice. Sholl analysis of different hippocampal areas showed a significant simplification of neuronal geometry in CA1 neurons of RNF10 KO compared to control animals (**Figure 5G, H**). Interestingly, we observed no difference in dendritic arborization of dentate gyrus neurons, suggesting a cell-specific effect of RNF10 deficiency within the hippocampus (**Figure 5I, J**).

### Silencing of RNF10 in the dCA1 leads to NMDAR-related molecular alterations

To understand the molecular mechanisms underlying the structural, functional and behavioral alterations induced by RNF10 silencing, we used hippocampal CA1 micro-dissected sections from RNF10-silenced and control mice to determine gene expression by RNA-seq. PCA analysis indicated (**Supplementary Figure 3A**) that samples were grouped very well according to their condition (shRNF10 vs. Scr), and it is possible to observe a difference in the clustering when controls are compared to shRNF10 samples (**Figure 6A**). The differential gene expression analysis led to identifying 78 upregulated genes and 367 downregulated genes in RNA samples upon comparing shRNF10 samples and Scr controls (**Figure 6A, B**). The significant differentially expressed genes are displayed in the Volcano plot (**Figure 6B**) and reported in **Supplementary Table 5**. The number of total normalized counts and total raw counts per sample obtained from the DESeq2 analysis are reported in **Supplementary Table 6**.

**Figure 6.**
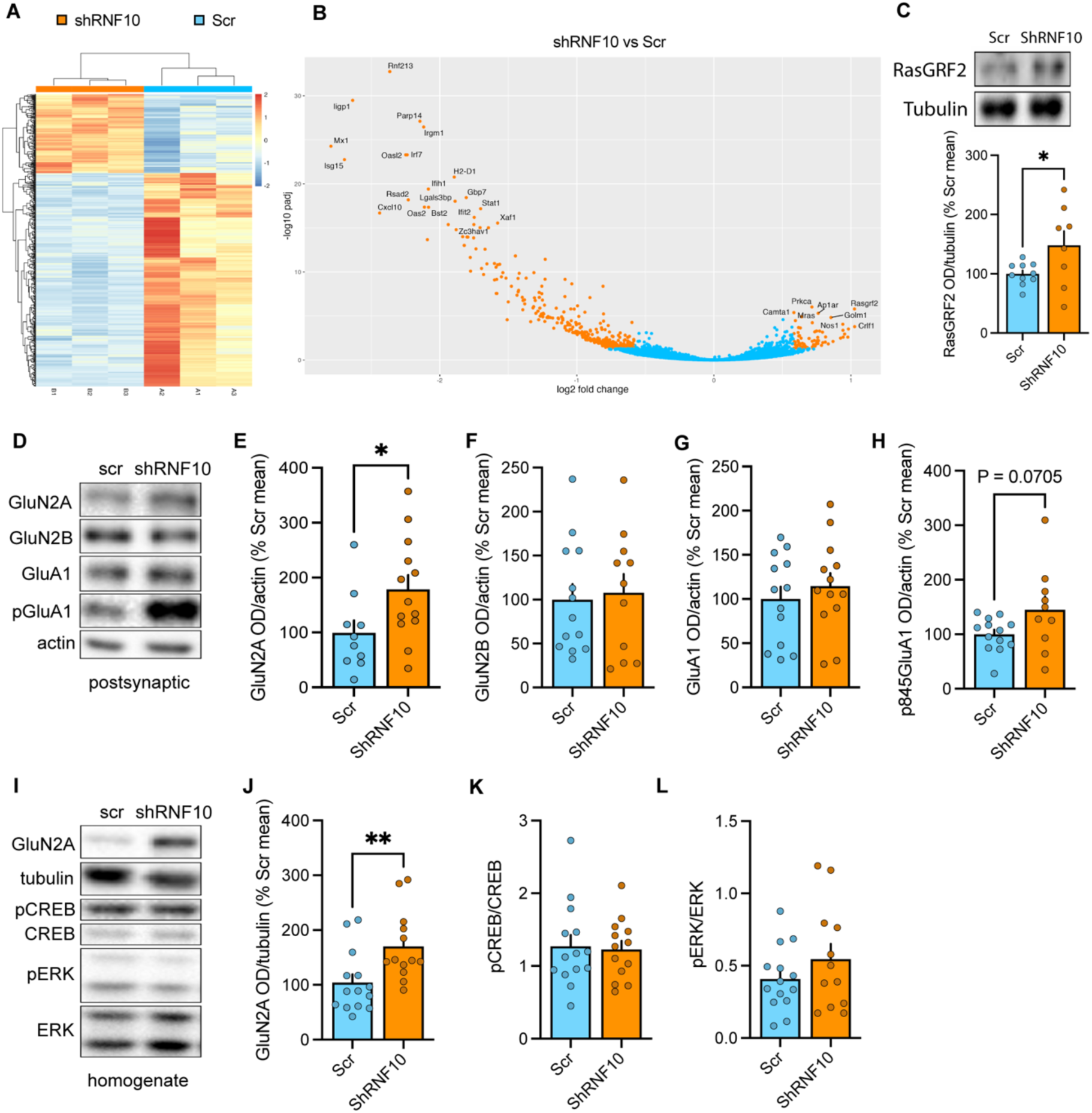
Silencing of RNF10 in the dCA1 leads to molecular alterations. (**A**) Heatmap showing the differentially expressed genes between ShRNF10 and Scr samples. The gene expression pattern is recurrent within replicates belonging to the same condition and is distinct when samples of different conditions are compared. (**B**) Volcano plot indicating the trend of the upregulated and downregulated genes in ShRNF10 compared to control Scr samples. (**C**) Western blot representative images (upper panel) and bar graph (lower panel) of densitometric quantification of RasGRF2 in total cell homogenate in Scr and ShRNF10 dCA1 samples (two-tailed unpaired *t-*test; RasGRF2: p = 0.0453, n = 10/8). The tubulin band was used for normalization. (**D**) Western blot representative images and bar graph of densitometric quantification of (**E**) GluN2A, (**F**) GluN2B, (**G**) GluA1 and (**H**) phosphorylated ser845 GluA1 subunit in triton-insoluble postsynaptic fractions (TIF) of Scr and ShRNF10 dCA1 samples (two-tailed unpaired *t-test*; GluN2A: p = 0.0308, n = 11/13; GluN2B: p = 0.7777, n = 13/11; GluA1; p = 0.4706, n = 13/group; p845GluA1: p = 0.0705, n = 13/10). The actin band was used for normalization. (**I**) Western blot representative images and bar graph of densitometric quantification in total cell homogenate of (**J**) GluN2A, (**K**) pCREB/CREB, and (**L**) pERK/ERK of Scr and ShRNF10 dCA1 samples (two-tailed unpaired *t*-test; GluN2A: p = 0.0082, n = 14/13; pCREB/CREB: p = 0.8306, n = 14/13; pERK/ERK: p = 0.2444, n = 14/12). For GluN2A, the tubulin band was used for normalization. pERK and pCREB proteins were normalized against ERK and CREB bands, respectively. *p < 0.05, **p < 0.01. Values are expressed as means ± s.e.m.

Many genes coding for synaptic proteins are included in the top 20 significantly downregulated and upregulated genes (**Supplementary Figure 3B**), confirming a role for RNF10 activity in regulating synaptic function. In particular, *Rasgrf2* (Ras Protein Specific Guanine Nucleotide Releasing Factor 2) was detected as one of the genes whose expression was most significantly upregulated by RNF10 silencing (**Figure 6B and Supplementary Figure 3B**). This is particularly relevant considering the role of RasGRF2 protein in the induction of long-term synaptic plasticity in hippocampal neurons (45, 46) and its function in coupling the activation of NMDARs to downstream signaling events (47, 48). Importantly, western blotting analysis further confirmed the increased expression of RasGRF2 protein in RNF10-silenced dCA1 compared to controls (**Figure 6C**).

RasGRF2 has been previously shown to be strictly coupled to the activation of synaptic-retained GluN2A-containing NMDARs (45, 46). Accordingly, we evaluated whether RNF10 silencing could also affect the localization of NMDAR subunits at postsynaptic sites subunit (**Figure 6D**). Western blotting analysis performed in a triton-insoluble postsynaptic fraction, enriched in postsynaptic density proteins, showed an aberrant increase in the GluN2A subunit in microdissected dCA1 areas from RNF10-silenced mice compared to controls without any alteration of the GluN2B subunit (**Figure 6E, F**). Analysis of GluN2A levels also showed a significant increase in the GluN2A subunit in the total homogenate (**Figure 6I, J**). RNF10 silencing in dCA1 did not modify the postsynaptic enrichment of the GluA1 subunit of AMPARs (**Figure 6G**) but induced an increase, although not statistically detected (p = 0.07), in the phosphorylation of the subunit at Ser845 (**Figure 6H**), which is known to be strictly related to receptor activation and induction of synaptic plasticity (49). Finally, analysis of two main signaling pathways downstream of ionotropic glutamate receptor activation, pCREB/CREB and pERK/ERK, did not show any alteration induced by RNF10 silencing, suggesting a specific alteration of RasGRF2 among NMDAR signaling proteins (**Figure 6K, L**).

Finally, to determine whether the molecular alterations identified following RNF10 silencing were specifically attributable to loss of RNF10, we next examined the expression of key candidates in the rescue condition (**Supplementary figure 4A**). While GluN2A levels were completely restored in ShRNF10 + ShResistant mice (**Supplementary figure 4B**), RasGRF2 shows just a partial recovery, with a trend toward normalization that did not reach control levels (**Supplementary figure 4C**). Overall, these findings suggest that restoring RNF10 expression in dCA1 partially rescues both behavioral and molecular alterations, while revealing a shift in strategy during reversal learning, highlighting the contribution of RNF10 signaling to hippocampal-dependent behavioral updating

## DISCUSSION

Cognitive flexibility is essential for adaptive behaviour and for maintaining brain health across the lifespan. When these mechanisms become dysregulated, as occurs during ageing and in several neurodevelopmental and neurodegenerative disorders, individuals often display behavioural rigidity and impaired adaptation to changes. Despite growing knowledge of the brain circuits that support flexible behaviour, the cellular and molecular mechanisms that translate neuronal activity into behavioural adaptation remain poorly understood. Here, we show that RNF10, known as an E3 ubiquitin ligase, also functions as a NMDAR-associated protein, and that this neuronal activity is relevant for cognitive flexibility. In detail, we show that the RNF10-mediated signaling pathway, linking activation of GluN2A-containing NMDARs to nuclear gene expression in the dCA1, is necessary for cognitive flexibility. Notably, the downregulation of RNF10, through gene deletion and silencing, leads to modifications of dendritic branching and spine morphology *in vivo*, specifically in the CA1. These morphological events are related to abnormal CA1 neuronal excitability, long-term plasticity, and changes in the expression of long-term synaptic plasticity markers. At the behavioral level, this dysregulated context leads to impaired ability to modify action-outcome associations in a dynamic environment, which is a fundamental aspect of cognitive flexibility. Thus, our results implicate RNF10 as a key factor *in vivo*, enabling the correct expression of cognitive flexibility.

Classical models of cognitive flexibility primarily implicate frontal cortical networks—including the dorsolateral and orbitofrontal prefrontal cortex and their corticostriatal loops—as key substrates for adapting behaviour to changing stimulus–outcome associations (50, 51), with relatively little emphasis on hippocampal contributions (3). However, some studies have proposed that the hippocampus could be central in the modulation of goal-directed behavior (8–10). consistent with these reports, here we showed that dCA1 is critical in mice to express cognitive flexibility in different behavioral tasks. Furthermore, we identified a molecular pathway linking GluN2A-containing NMDARs to specific transcriptional programs implicated in these behavioral functions. In particular, we found that NMDAR-dependent synapse-to-nucleus communication mediated by RNF10 is required in the dCA1 for intact reversal but not initial learning, probably because of persistence in perseverative errors. A recent study has shown that CA2, which sends its projections to the CA1, contributes to early reversal learning in a Morris water maze (10). Here, using an operant visual cue response discrimination, we extend the role of the hippocampus in flexible behavior to a non-navigational context. In this task, RNF10 might be necessary for the dCA1 to update choice–outcome relations when the cognitive load becomes more relevant, such as upon changes in reward contingencies during reversal learning (**Figure 7**). This would be consistent with the role of the hippocampus as a ‘comparator’, in synergy with the prefrontal cortex, of expected action outcomes to actual action outcomes (9). The dCA1 detects novelty by comparing expected events derived from prior memories with actual current events (8). This operation results in memory strengthening or flexible reformulation of behavior and memory updating (**Figure 7**). Consistent with this scenario, we have shown that RNF10 silencing hampers the long-term attenuation of aversive contextual memory, suggesting dysfunctional updating of acquired memories under novel contingencies.

**Figure 7.**
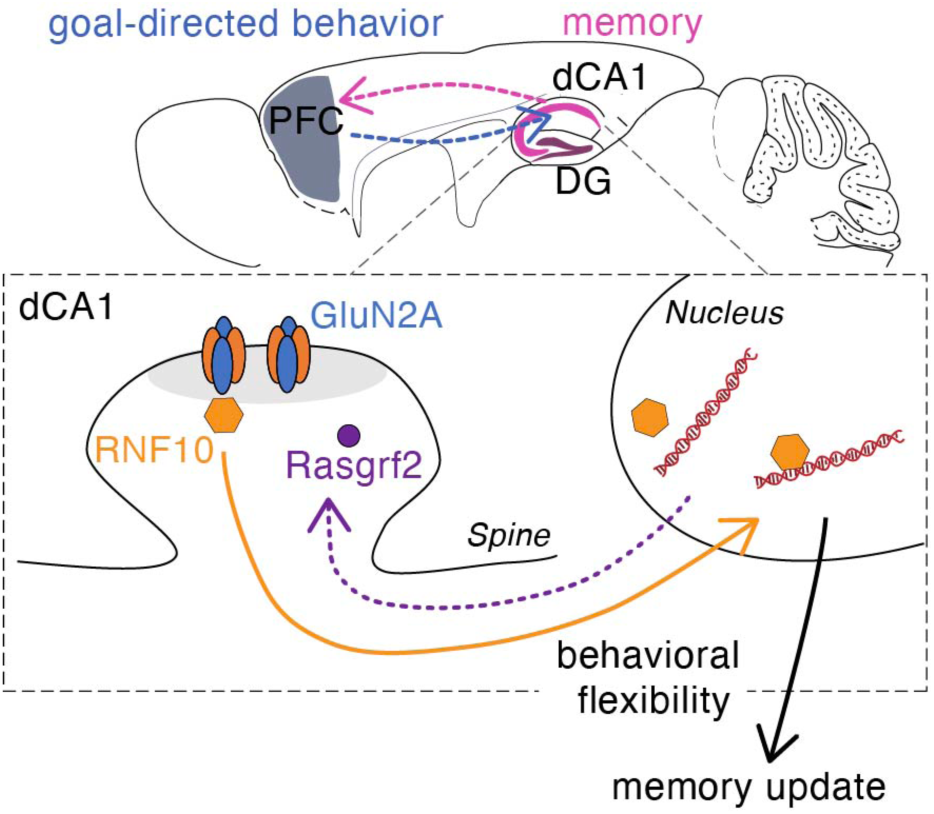
A model of the role of RNF10 in cognitive flexibility. In this model, the dCA1 would be required to update choice–outcome relations in goal-directed behavior. Activation of dCA1 neurons through stimulation of NMDA receptors triggers RNF10 signalling to the nucleus to modulate gene expression of synaptic proteins. Among these genes, RasGRF2 is critically involved in the induction of long-term synaptic plasticity. This process allows plastic rearrangements of dendritic spines and connections to modified and update stored short-term memories into new ones.

This cognitive phenotype was confirmed by molecular alterations: an RNAseq assay and subsequent biochemical validation identified the Ras-GRF2 calcium sensor as one of the gene products most affected by RNF10 silencing in dCA1 neurons. Interestingly, Ras-GRF2 has been previously characterized as a key element in the signaling cascade downstream LTP induction mediated by activation of GluN2A-containing NMDARs in CA1 pyramidal neurons (45, 46). This is highly relevant considering that (i) RNF10 interacts specifically with GluN2A-containing NMDARs and does not bind GluN2B and (ii) its nuclear translocation is blocked by antagonists selective for the GluN2A but not the GluN2B subunit (26). On this view, the observed alteration of GluN2A but not GluN2B protein levels in CA1 homogenates from mice transduced with the viral vector expressing RNF10 shRNA supports the specific correlation between RNF10 activity and GluN2A-containing NMDARs. These results also confirm a putative role for GluN2A as a player in cognitive flexibility, as recently suggested by a study showing that GluN2A pharmacological blockade impairs reversal learning in mice (52).

We previously found that RNF10 downregulation *in vitro* in primary hippocampal neurons was associated with a decreased number of dendritic spines (26) and a significant simplification of dendritic arborization (27), suggesting that RNF10 might have a global effect on dendritic architecture. Here, in RNF10 KO mice, we confirmed that RNF10 is needed for a correct formation of the dendritic tree in CA1 but not in DG neurons. However, CA1 pyramidal neurons from both mice transduced with the shRNA RNF10 construct and RNF10 KO mice did not result in a reduction in spine density but in a significant alteration in the spine morphology with a reduction in spine head width and length. Overall, these data, together with neuroanatomical observations, also confirm the structural role of RNF10 in regulating neuronal morphology in an *in vivo* setting. Interestingly, these morphological alterations are paralleled by significant impairment of long-term plasticity in RNF10 CA1 neurons associated with divergent priming effects of bursts of high-frequency stimulations in RNF10 KO compared to control neurons. Conversely, synaptic transmission was not altered in Schaffer collaterals to the CA1 synapse of RNF10 KO mice. In agreement with these data, we previously showed a deficit in the long-lasting increase in the amplitude and frequency of mEPSCs following the induction of chemical LTP in the absence of RNF10 in primary hippocampal neurons (26).

In conclusion, we highlighted the role of NMDAR-dependent transcriptional regulation in dorsal CA1, as mediated by RNF10, as a key event to maintain cognitive flexibility.

## Acknowledgments

We thank Marta Ornaghi for her technical assistance. This work was supported by PNRR, Missione 4 – Componente 2– Investimento 1.3, finanziato dall’Unione europea – NextGenerationEU “Fascination” to M.D. and FIS00000560 – “Stone” to M.D, the Ministry of University and Research (PRIN20202THZAW to MDL, PSR2023 and 2025 – Azione B – linea 2 to DS) and to the Department of Pharmacological and Biomolecular Sciences “Rodolfo Paoletti,” University of Milan (MUR - Progetto Eccellenza 2023–2027) and the Italian Ministry of Health (Ricerca Corrente and 5 × 1.000 funds) (to NM).

The authors declare that they have no competing interests.

## Author contributions

Conceptualization: ER, NC, EM, FG, MDL, DS; Methodology: ER, NC, MI, FLG, FG, SCC, MS, NM; Investigation: ER, NC, AR, MI, FLG, LD, EZ, LP, GB; Analysis: ER, NC, FG, GB, DS; Funding acquisition: DS, MDL, NM; Project administration: FG, MDL, DS; Supervision: NM, MS, CM, BY, EM, FG, MDL, DS; Writing – original draft: ER, NC, FG, SCC, BY, FG, DS; Writing – review & editing: CM, EM, FG, MDL, DS.

## Data availability

All raw data concerning RNA-seq have been deposited in NCBI’s Gene Expression Omnibus (53) and are accessible through GEO Series accession number 245221 at https://www.ncbi.nlm.nih.gov/geo/query/acc.cgi?acc=GSE245221.

