## Supplemental information for "Hippocampal Ring Finger Protein 10-dependent signaling supports cognitive flexibility"

### Supplementary Materials

#### Supplementary Figure 1

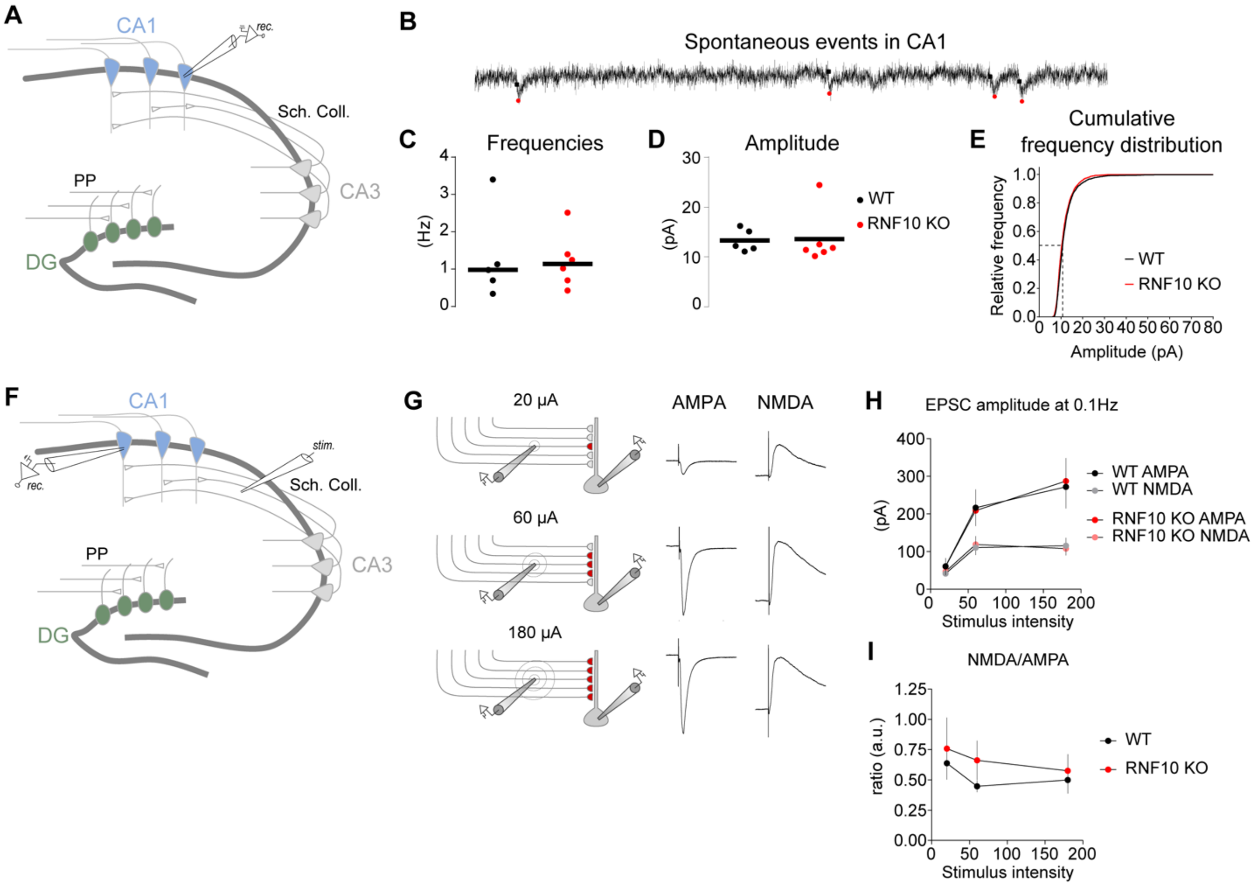


**Supplementary Figure 1. The absence of RNF10 does not alter synaptic transmission in Schaffer collaterals to CA1 synapses.** (**A**) Schematic of the main hippocampal sub-regions (dentate gyrus: DG, CA3, CA1). (**B-E**) Results obtained by recording spontaneous synaptic activities in CA1 pyramidal neurons. (**B**) Electrophysiological recording representing spontaneous synaptic events in CA1. (**C, D**) The frequency and amplitude of the spontaneous events were not altered by the absence of RNF10 (C; Mann–Whitney test, Sum of ranks = 34/32, U = 11, n = 5/6, p = 0.6991, D; Mann–Whitney test, Sum of ranks = 34/32, U = 11, n = 5/6, p = 0.5368). (**E**) Cumulative frequencies of EPSC amplitude in WT or RNF10 KO mice (Kolmogorov–Smirnov test, n = 1936/2153, p = 0.0003). (**F**) Schematic of the stimulation of the Schaffer collaterals while synaptic responses are recorded in CA1 pyramidal neurons as in ‘G’ to ‘K’. (**G**) Schematic showing the stimulation of an increasing number of collateral fibers by increasing the intensity of the stimulus. Representative AMPA and NMDA responses are shown. (**H**) Graphs of the amplitude of the EPSCs of AMPAR- or NMDAR-mediated currents (Two-way ANOVA, AMPA: stimulus intensity, $F_{(2, 57)}$ = 12.79, p <0.0001; Genotype, F (1, 57) = 0.004052, p = 0.9495; interaction, F (2, 57) = 0.03530, p = 0.9653. NMDA: Stimulus Intensity, $F_{(2, 57)}$ = 10.38, p = 0.0001; genotype, F (1, 57) = 0.01807, p = 0.8935; interaction,

$F_{(2, 57)} = 0.1157$, $p = 0.8910$. (**I**) Ratio of NMDAR responses normalized to AMPAR responses (Two-way ANOVA, stimulus intensity, $F = 0.7602$, $p = 0.4731$; Genotype, $F_{(1, 48)} = 1.461$, $p = 0.2326$; interaction, $F_{(2, 48)} = 0.1338$, $p = 0.8751$). Values are expressed as means ± s.e.m.

**Supplementary Figure 2**

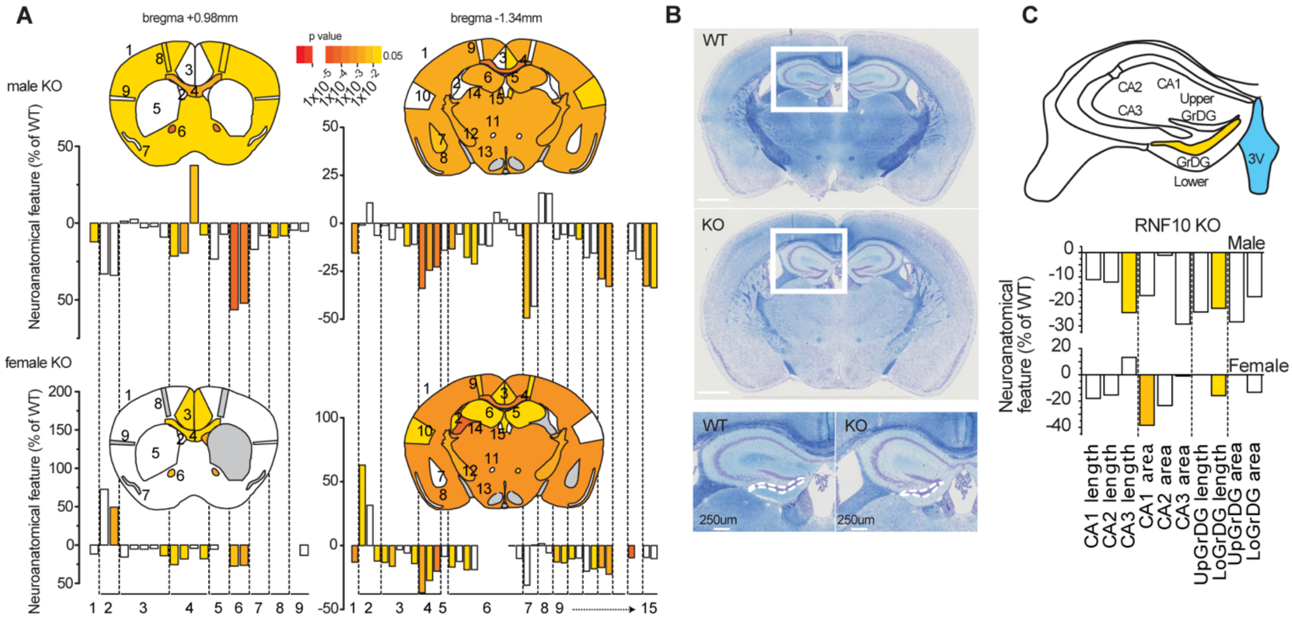


**Supplementary Figure 2. Neuroanatomical structural abnormalities of hippocampus in RNF10 KO mice.** (**A**) Schematic representation of the brain regions analyzed in adult male (top) and female (bottom) mice (16 weeks old) using a quantitative 2D histological approach. A total of 63 morphological parameters were assessed across 24 brain regions (**Supplementary Table 1**) by systematically quantifying two coronal planes located at Bregma +0.98 mm and Bregma −1.34 mm (**Supplementary Table 2**). The color map indicates parameters reaching statistical significance ($p < 0.05$), while gray regions correspond to areas where the number of observations was insufficient to perform statistical analysis. Histograms represent the fold increase or decrease expressed as a percentage relative to wild-type (WT) mice. Dashed lines indicate regrouped parameters within each brain region. The analysis revealed a consistent microcephaly phenotype in RNF10 KO mice of both sexes. At Bregma −1.34 mm, the total brain area was reduced by 15% in males ($P = 0.0023$) and 13% in females ($P = 0.00072$) compared with WT controls. This reduction was accompanied by a decrease in dorsal hippocampus size of 13% in males ($P = 0.0013$) and 16% in females ($P = 0.044$). Several commissural structures—including the genu and body of the corpus callosum, the anterior commissure, the internal capsule, and the dorsal hippocampal commissure—were also significantly reduced in RNF10 KO mice (Supplementary Table 3). For example, the corpus callosum body area decreased by 34% in males ($P = 0.00029$) and 36% in females ($P = 0.0014$). (**B**) Representative Nissl–

Luxol double-stained coronal brain sections from WT (top) and RNF10 KO (bottom) mice illustrating the reduction in overall brain size and the abnormal morphology of the hippocampus (white frame). The dashed line marks the lower arm of the granular layer of the dentate gyrus, which showed the most pronounced structural alterations. (**C**) Finer-scale characterization of hippocampal alterations in RNF10 KO mice. Top: schematic representation of the dorsal hippocampus at Bregma −1.34 mm. Bottom: histograms showing the areas of individual hippocampal layers quantified through a detailed morphometric analysis (Supplementary Table 4). The hippocampal phenotype was comparable between sexes, with a significant reduction in the length of the lower arm of the dentate gyrus granular layer (GrDG low), decreased by 18% in males (P = 0.038) and 17% in females (P = 0.016) relative to WT mice. Abbreviations: CA1 = layer 1 of the cornus ammonis; CA2/3 = layers 2 and 3 of the cornus ammonis; GrDG up = upper arm of the granular layer of the dentate gyrus; GrDG low = lower arm of the granular layer of the dentate gyrus. Overall, this systematic neuroanatomical analysis indicates that RNF10 depletion leads to a robust phenotype characterized by microcephaly, hippocampal hypoplasia, and commissural abnormalities, with no major sex-specific differences.

**Supplementary Figure 3**

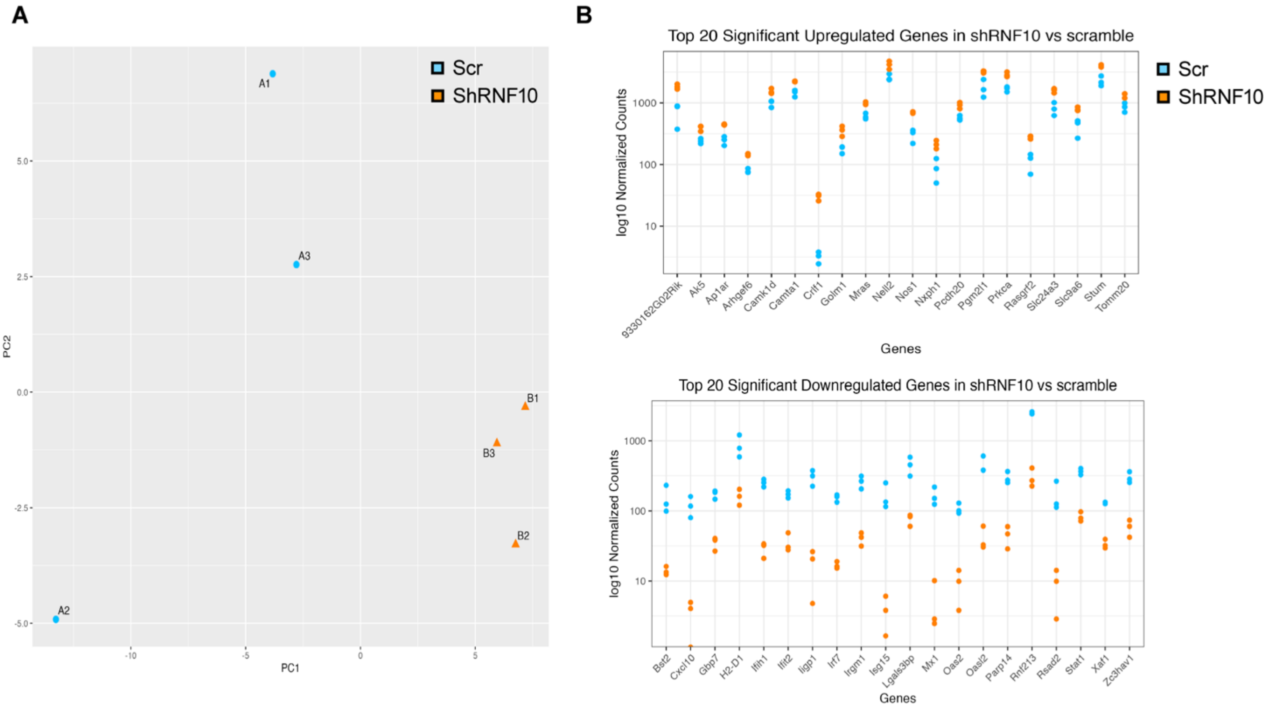


**Supplementary Figure 3. Gene expression effect induced by silencing RNF10 in the dCA1 of mice**. (**a**) Principal component analysis of transformed reads. Samples cluster according to their experimental condition. (**b**) Dot plot of the normalized counts showing the differential expression of the top 20 upregulated and downregulated genes in the shRNA RNF10 and scr control for comparison.

**Supplementary Figure 4**

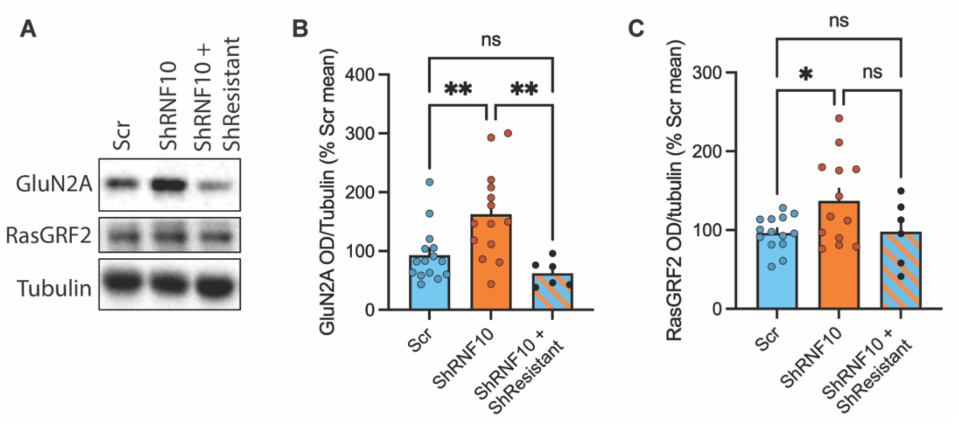


**Supplementary Figure 4. Rescue of cognitive proteins expression in ShRNF10+ShResistant mice.** (**A**) Western blot representative images and bar graph of densitometric quantification of (**B**) GluN2A (One-way ANOVA with Tukey's post hoc test; Scr vs ShRNF10 p=0.0076; Scr vs Sh+ShResistant p=0.5268; ShRNF10 vs Sh+ShResistant p=0.0033) and (**C**) RasGRF2 (One-way ANOVA with Tukey's post hoc test; Scr vs ShRNF10 p=0.0408; Scr vs Sh+ShResistant p=0.9969; ShRNF10 vs Sh+ShResistant p=0.1503) in total homogenate of Scr (n=15), ShRNF10 (n=14) and ShRNF10 + ShResistant (n=6) dCA1 samples. The tubulin band was used for normalization. Values are expressed as means ± s.e.m. *p < 0.05, **p < 0.01, ***p < 0.001, ****p<0.0001.

**Supplementary Table 1** List of 63 neuroanatomical measurements. This table provides morphological phenotypes studied in the coronal plane. Stereotaxic coordinates (column A) of the two coronal sections are indicated. Association number of each parameter with a brain region is indicated in column B. The abbreviations, the description of each parameter and the unit of the measurement are given in columns C, D and E, respectively.

| Section | Region_ID | Parameters | Description | Units |
|---|---|---|---|---|
| Bregma +0.98 mm | 1 | 1_TBA | Total Brain Area | cm² |
| | 2 | 1_LV_L | Area of Lateral Ventricule_Left | cm² |
| | | 1_LV_R | Area of Lateral Ventricule_Right | cm² |
| | 3 | 1_Cg_L | Area of Cingulate gyrus_Left | cm² |
| | | 1_Cg_R | Area of Cingulate gyrus_Right | cm² |
| | | 1_Cg_Width_L | Width of Cingulate gyrus_Left | cm |
| | | 1_Cg_Width_R | Width of Cingulate gyrus_Right | cm |
| | | 1_Cg_Height | Height of Cingulate gyrus | cm |
| | 4 | 1_gcc | Area of Genu of corpus callosum | cm² |
| | | 1_gcc_Width_T | Width of genu of corpus callosum_Top | cm |
| | | 1_gcc_Width_B | Width of genu of corpus callosum_Bottom | cm |
| | | 1_gcc_Height | Height of genu of corpus callosum | cm |
| | 5 | 1_CPu_L | Area of Caudate Putamen_Left | cm² |
| | | 1_CPu_R | Area of Caudate Putamen_Right | cm² |
| | 6 | 1_aca_L | Area of anterior commissure_Left | cm² |
| | | 1_aca_R | Area of anterior commissure_Right | cm² |
| | 7 | 1_Pir_L | Area of Piriform cortex_Left | cm² |
| | | 1_Pir_R | Area of Piriform cortex_Right | cm² |
| | 8 | 1_M1_L | Height of primary Motor cortex_Left | cm |
| | | 1_M1_R | Height of primary Motor cortex_Right | cm |
| | 9 | 1_S2_L | Height of secundary somatosensory cortex_Left | cm |
| | | 1_S2_R | Height of secundary somatosensory cortex_Right | cm |
| Bregma -1.34 mm | 1 | 2_TBA | Total Brain Area | cm² |
| | 2 | 2_LV_L | Area of Lateral ventricule_Left | cm² |
| | | 2_LV_R | Area of Lateral Ventricule_Right | cm² |
| | | 2_D3V | Area of dorsal third ventricule | cm² |
| | 3 | 2_RSGc_L | Area of Retrosplenial Granular cortex_Left | cm² |
| | | 2_RSGc_R | Area of Retrosplenial Granular cortex_Right | cm² |
| | | 2_RSGc_Width_L | Width of Retrosplenial Granular cortex_Left | cm |
| | | 2_RSGc_Width_R | Width of Retrosplenial Granular cortex_Right | cm |
| | | 2_RSGc_Height | Height of Retrosplenial Granular cortex | cm |
| | 4 | 2_cc | Area of corpus callosum | cm² |
| | | 2_cc_Width | Width of corpus callosum | cm |

| | | | | |
|---|---|---|---|---|
| | | 2_cc_Height | Height of corpus callosum | cm |
| | 5 | 2_dhc | Area of hippocampal commissure | cm² |
| | 6 | 2_HP | Area of Hippocampus | cm² |
| | | 2_TILpy | Total Internal Length of pyramidal layer | cm |
| | | 2_DG_L | Length of Dentate Gyrus_Left | cm |
| | | 2_DG_R | Length of Dentate Gyrus_Right | cm |
| | | 2_Mol_L | Length of Molecular Layer_Left | cm |
| | | 2_Mol_R | Length of Molecular Layer_Right | cm |
| | | 2_Rad_L | Length of Radial Layer_Left | cm |
| | | 2_Rad_R | Length of Radial Layer_Right | cm |
| | | 2_Or_L | Length of Orion Layer_Left | cm |
| | | 2_Or_R | Length of Orion Layer_Right | cm |
| | 7 | 2_AM_L | Area of Amygdala_Left | cm² |
| | | 2_AM_R | Area of Amygdala_Right | cm² |
| | 8 | 2_Pir_L | Area of Piriform cortex_Left | cm² |
| | | 2_Pir_R | Area of Piriform cortex_Right | cm² |
| | 9 | 2_M1_L | Height of primary Motor cortex_Left | cm |
| | | 2_M1_R | Height of primary Motor cortex_Right | cm |
| | 10 | 2_S2_L | Height of secundary Somatosensory cortex_Left | cm |
| | | 2_S2_R | Height of secundary Somatosensory cortex_Right | cm |
| | 11 | 2_mt_L | Area of mammillothalamic tract_Left | cm² |
| | | 2_mt_R | Area of mammillothalamic tract_Right | cm² |
| | 12 | 2_ic_L | Area of internal capsule_Left | cm² |
| | | 2_ic_R | Area of internal capsule_Right | cm² |
| | 13 | 2_opt_L | Area of optic tract_Left | cm² |
| | | 2_opt_R | Area of optic tract_Right | cm² |
| | 14 | 2_fi_L | Area of fimbria of hippocampus_Left | cm² |
| | | 2_fi_R | Area of fimbria of hippocampus_Right | cm² |
| | 15 | 2_Hb_L | Area of habenular nucleus_Left | cm² |
| | | 2_Hb_R | Area of habenular nucleus_Right | cm² |

**Supplementary Table 2** List of 10 additional neuroanatomical measurements pertaining to the dorsal hippocampus.

| Section | Region_ID | Parameters | Description | Units |
|---|---|---|---|---|
| Bregma -1.34 mm | 1 | CA1_area | Area of the CA1 layer of the hippocampus | $cm^2$ |
| | | CA1_length | Length of the CA1 layer of the hippocampus | cm |
| | 2 | CA2_area | Area of the CA2 layer of the hippocampus | $cm^2$ |
| | | CA2_length | Length of the CA2 layer of the hippocampus | cm |
| | 3 | CA3_area | Area of the CA3 layer of the hippocampus | $cm^2$ |
| | | CA3_length | Length of the CA3 layer of the hippocampus | cm |
| | 4 | GrDG_upper_area | Area of the granular upper layer of the dentate gyrus | $cm^2$ |
| | | GrDG_upper_length | Length of the granular upper layer of the dentate gyrus | cm |
| | 5 | GrDG_lower_area | Area of the granular lower layer of the dentate gyrus | $cm^2$ |
| | | GrDG_lower_length | Length of the granular lower layer of the dentate gyrus | cm |

**Supplementary Table 3** Raw neuroanatomical measurements for male and female KO mice

| Sex | Male | Male | Male | Male | Male | Male | Male | Male | Male | Male | Male | Male | Female | Female | Female | Female | Female | Female |
|---|---|---|---|---|---|---|---|---|---|---|---|---|---|---|---|---|---|---|
| **Barcode** | M018663 | M018663 | M01868 | M01869 | M01869 | M01874 | M018796 | M01889 | M018895 | M01869 | M01874 | M0189 | A000013 | A000013 | A00001 | A000013 | A000013 | A00001396 |
| **Gene** | Rnf10 | Rnf10 | Rnf10 | Rnf10 | Rnf10 | Rnf10 | Rnf10 | Rnf10 | Rnf10 | Rnf10 | Rnf10 | Rnf10 | Rnf10 | Rnf10 | Rnf10 | Rnf10 | Rnf10 | Rnf10 |
| **Genotype** | WT | WT | WT | WT | WT | WT | WT | WT | WT | HOM | HOM | HOM | WT | WT | WT | HOM | HOM | HOM |
| **1_TBA** | 0,27888 | 0,3382 | 0,3324 | 0,292 | 0,34 | 0,3202 | 0,2786 | 0,294 | 0,2949 | 0,281 | 0,2784 | 0,251 | 0,3542 | 0,2978 | 0,3022 | 0,28138 | 0,2807 | |
| **1_LV_L** | 0,00116 | 0,00158 | 0,0048 | 0,003 | 7E-04 | 0,0029 | 0,001 | 0,002 | 0,002 | 0,0023 | 0,001 | 0,001 | 0,001 | 0,0011 | 0,0015 | 0,00118 | 0,0026 | 0,0026 |
| **1_LV_R** | 0,00227 | 0,00088 | 0,0045 | 0,004 | 0,001 | 0,0034 | 0,0011 | 0,001 | 0,0014 | 0,0015 | 0,0018 | 0,001 | 0,0011 | 0,0014 | 0,0012 | 0,00172 | 0,0018 | 0,0019 |
| **1_Cg_L** | 0,0083 | 0,00984 | 0,0092 | 0,008 | 0,01 | 0,0089 | 0,008 | 0,008 | 0,0085 | 0,0081 | 0,0096 | | 0,0098 | 0,009 | 0,0084 | 0,00791 | 0,0074 | |
| **1_Cg_R** | 0,00776 | 0,01057 | 0,0094 | 0,009 | 0,01 | 0,0103 | 0,0082 | 0,009 | 0,0084 | 0,0086 | 0,0101 | | 0,0097 | 0,0079 | 0,0089 | 0,00835 | 0,0084 | |
| **1_Cg_Width_L** | 0,08122 | 0,09514 | 0,0919 | 0,086 | 0,097 | 0,0866 | 0,0865 | 0,08 | 0,0809 | 0,0836 | 0,0864 | 0,084 | 0,0904 | 0,0864 | 0,0889 | 0,08856 | 0,0816 | 0,0809 |
| **1_Cg_Width_R** | 0,08582 | 0,09868 | 0,0919 | 0,082 | 0,095 | 0,0923 | 0,0924 | 0,093 | 0,0955 | 0,0871 | 0,0936 | 0,088 | 0,0904 | 0,0787 | 0,0889 | 0,08637 | 0,0787 | 0,0809 |
| **1_Cg_Height** | 0,14271 | 0,15491 | 0,152 | 0,153 | 0,171 | 0,1596 | 0,1465 | 0,145 | 0,141 | 0,1303 | 0,1607 | 0,123 | 0,1501 | 0,1563 | 0,1458 | 0,12792 | 0,1312 | |
| **1_gcc** | 0,00818 | 0,0107 | 0,0094 | 0,006 | 0,01 | 0,0095 | 0,0097 | 0,009 | 0,0089 | 0,0079 | 0,0062 | 0,007 | 0,0078 | 0,0069 | 0,0073 | 0,00568 | 0,0052 | |
| **1_gcc_Height** | 0,04237 | 0,0445 | 0,0416 | 0,036 | 0,045 | 0,0448 | 0,0425 | 0,04 | 0,0452 | 0,0346 | 0,0311 | 0,037 | 0,0328 | 0,0361 | 0,0335 | 0,02915 | 0,0268 | |
| **1_gcc_Width_B** | 0,05366 | 0,08092 | 0,055 | 0,048 | 0,07 | 0,0717 | 0,0626 | 0,078 | 0,0795 | 0,0859 | 0,093 | 0,096 | 0,07 | 0,0547 | 0,0729 | 0,06778 | 0,0583 | |
| **1_gcc_Width_T** | 0,15331 | 0,16876 | 0,1531 | 0,144 | 0,151 | 0,1568 | 0,1578 | 0,155 | 0,1509 | 0,1545 | 0,1355 | 0,138 | 0,1388 | 0,1487 | 0,1516 | 0,12792 | 0,1122 | |
| **1_CPu_L** | 0,02118 | 0,04419 | 0,0362 | 0,033 | 0,037 | 0,0338 | 0,0307 | 0,036 | 0,0359 | 0,0229 | 0,0268 | 0,029 | 0,0293 | 0,0261 | 0,03 | | 0,0278 | 0,026 |
| **1_CPu_R** | 0,02772 | 0,03379 | 0,034 | 0,032 | 0,042 | 0,0326 | 0,0269 | 0,03 | 0,0304 | 0,0321 | 0,0329 | 0,024 | 0,0366 | 0,0316 | 0,0307 | | | 0,0255 |
| **1_aca_L** | 0,00055 | 0,00075 | 0,0007 | 6E-04 | 7E-04 | 0,0006 | 0,0007 | 8E-04 | 0,0008 | 0,0003 | 0,0003 | 2E-04 | 0,0006 | 0,0005 | 0,0006 | 0,00044 | 0,0004 | 0,0004 |
| **1_aca_R** | 0,00058 | 0,00079 | 0,0007 | 5E-04 | 7E-04 | 0,0006 | 0,0007 | 7E-04 | 0,0008 | 0,0003 | 0,0004 | 3E-04 | 0,0006 | 0,0006 | 0,0006 | 0,00041 | 0,0004 | 0,0004 |
| **1_Pir_L** | 0,00125 | 0,00239 | 0,0018 | 0,002 | 0,002 | 0,0014 | 0,0014 | 0,002 | 0,0014 | 0,0012 | 0,0017 | 0,001 | 0,0015 | 0,0014 | 0,0015 | | 0,0013 | |
| **1_Pir_R** | 0,00121 | 0,00235 | 0,0017 | 0,001 | 0,002 | 0,0014 | 0,0014 | 0,001 | 0,0014 | 0,0015 | 0,0019 | 0,001 | 0,0017 | 0,0013 | 0,0014 | 0,00139 | | |
| **1_M1_L** | 0,11868 | 0,12121 | 0,1254 | 0,127 | 0,128 | 0,1257 | 0,1129 | 0,12 | 0,1179 | 0,1159 | | 0,105 | 0,12 | 0,1213 | 0,1131 | 0,10395 | | |
| **1_M1_R** | 0,1159 | 0,11888 | 0,1237 | 0,123 | 0,125 | 0,1356 | 0,1133 | 0,12 | 0,1199 | 0,1174 | 0,1128 | 0,104 | 0,1218 | 0,1129 | 0,1192 | 0,10129 | | |
| **1_S2_L** | 0,10351 | 0,10279 | 0,1131 | 0,095 | 0,102 | 0,1004 | 0,105 | 0,098 | 0,0956 | 0,1126 | 0,096 | 0,083 | | 0,1082 | | 0,08746 | 0,0977 | |
| **1_S2_R** | 0,09659 | 0,10223 | 0,1086 | 0,097 | 0,105 | 0,1038 | 0,0923 | 0,094 | 0,0926 | 0,101 | | 0,087 | 0,105 | 0,0929 | 0,0977 | 0,08746 | | 0,0831 |
| **2_TBA** | 0,41148 | 0,47674 | 0,4557 | 0,39 | 0,461 | 0,449 | 0,461 | 0,435 | 0,4316 | 0,3886 | 0,3573 | 0,372 | 0,4441 | 0,4447 | 0,4574 | 0,38618 | 0,3996 | 0,3867 |
| **2_LV_L** | 0,00233 | 0,0008 | 0,0071 | 0,004 | 0,002 | 0,0039 | 0,0013 | 8E-04 | 0,003 | 0,0023 | 0,0027 | 0,003 | 0,0028 | 0,0027 | 0,0018 | 0,00417 | 0,0034 | 0,0043 |
| **2_LV_R** | 0,00141 | 0,00114 | 0,0046 | 0,005 | 9E-04 | 0,002 | 0,0012 | 8E-04 | 0,0032 | 0,0033 | 0,0021 | 0,002 | 0,0024 | 0,003 | 0,0026 | 0,00272 | 0,0038 | 0,0041 |
| **2_D3V** | 0,00205 | 0,00218 | 0,0033 | 0,002 | 0,003 | 0,0031 | 0,0019 | | 0,0032 | 0,0028 | 0,0018 | 0,003 | 0,0014 | 0,0014 | 0,0015 | 0,00127 | 0,0012 | 0,0013 |
| **2_RSGc_L** | 0,00412 | 0,00469 | 0,0046 | 0,004 | 0,005 | 0,0058 | 0,0046 | 0,005 | 0,0038 | 0,0042 | 0,0056 | 0,004 | 0,0052 | 0,0053 | 0,0047 | 0,00454 | 0,0043 | 0,0044 |
| **2_RSGc_R** | 0,00424 | 0,00476 | 0,0046 | 0,005 | 0,005 | 0,0048 | 0,0053 | 0,005 | 0,0038 | 0,0042 | 0,0052 | 0,003 | 0,0054 | 0,0054 | 0,0051 | 0,00419 | 0,0044 | 0,0047 |
| **2_RSGc_Width_L** | 0,06478 | 0,06597 | 0,0683 | 0,066 | 0,069 | 0,0738 | 0,0658 | 0,069 | 0,0641 | 0,0625 | 0,0641 | 0,071 | 0,07 | 0,0711 | 0,0656 | 0,06969 | 0,0645 | 0,0656 |
| **2_RSGc_Width_R** | 0,06561 | 0,07071 | 0,0678 | 0,067 | 0,074 | 0,0761 | 0,0741 | 0,079 | 0,07 | 0,0653 | 0,0653 | 0,059 | 0,07 | 0,0678 | 0,07 | 0,06669 | 0,0612 | 0,0678 |
| **2_RSGc_Height** | 0,0883 | 0,09805 | 0,0913 | 0,091 | 0,109 | 0,1014 | 0,0929 | 0,098 | 0,0846 | 0,0924 | | 0,076 | 0,0995 | 0,1137 | 0,106 | 0,08637 | 0,0918 | 0,0962 |
| **2_cc** | 0,00369 | 0,00296 | 0,0036 | 0,003 | 0,003 | 0,0037 | 0,0033 | 0,003 | 0,0036 | 0,0026 | 0,0017 | 0,002 | 0,0032 | 0,0028 | 0,0031 | 0,00179 | 0,0019 | 0,0021 |
| **2_cc_Height** | 0,03062 | 0,02372 | 0,0263 | 0,028 | 0,025 | 0,0299 | 0,0304 | 0,026 | 0,0285 | 0,0231 | 0,0188 | 0,021 | 0,0262 | 0,0251 | 0,0208 | 0,01749 | 0,0186 | 0,0164 |
| **2_cc_Width** | 0,10881 | 0,11335 | 0,1009 | 0,106 | 0,109 | 0,1164 | 0,1097 | 0,104 | 0,0995 | 0,0918 | 0,0744 | 0,083 | 0,1126 | 0,1093 | 0,1071 | 0,08746 | 0,0864 | 0,0892 |
| **2_dhc** | 0,00032 | 0,00037 | 0,0004 | 4E-04 | 3E-04 | 0,0005 | 0,0005 | 4E-04 | 0,0005 | 0,0004 | 0,0003 | 3E-04 | 0,0002 | 0,0002 | 0,0003 | 0,00027 | 0,0002 | 0,0002 |
| **2_HP** | 0,03198 | 0,03607 | 0,0369 | 0,033 | 0,033 | 0,0361 | 0,0336 | 0,037 | 0,036 | 0,0305 | 0,0306 | 0,029 | 0,029 | 0,0247 | 0,0294 | 0,02385 | 0,0233 | 0,022 |
| **2_TILpy** | 0,53472 | 0,56542 | 0,5882 | 0,555 | 0,546 | 0,576 | 0,5358 | 0,599 | 0,6095 | 0,5563 | 0,5324 | 0,518 | 0,5186 | 0,4544 | 0,5429 | 0,44978 | 0,4372 | 0,4393 |
| **2_DG_L** | 0,14112 | 0,15701 | 0,1375 | 0,129 | 0,15 | 0,1499 | 0,1757 | 0,178 | 0,1771 | 0,1265 | 0,1317 | 0,124 | 0,1407 | | 0,1333 | 0,11196 | 0,105 | 0,1162 |
| **2_DG_R** | 0,13687 | 0,16764 | 0,1472 | 0,131 | 0,13 | 0,149 | 0,1281 | 0,164 | 0,1702 | 0,1049 | 0,1181 | 0,125 | 0,1585 | 0,1248 | 0,1177 | 0,1109 | 0,106 | |
| **2_Mol_L** | | 0,01015 | 0,0095 | 0,008 | | 0,0117 | 0,0126 | 0,008 | 0,0069 | 0,0088 | 0,0103 | 0,007 | | | | | | |
| **2_Mol_R** | | 0,00875 | 0,0084 | 0,007 | | 0,011 | 0,0095 | 0,011 | 0,0088 | 0,008 | 0,0073 | 0,009 | | | | | | |
| **2_Rad_L** | 0,02881 | 0,01894 | 0,0266 | 0,026 | 0,026 | 0,0279 | 0,0207 | 0,023 | 0,0219 | 0,0279 | 0,0258 | 0,024 | 0,0175 | | | 0,01859 | | |

| | | | | | | | | | | | | | | | | | | |
|---|---|---|---|---|---|---|---|---|---|---|---|---|---|---|---|---|---|---|
| **2_Rad_R** | 0,02151 | 0,02132 | 0,0274 | 0,028 | 0,021 | 0,023 | | 0,022 | 0,0241 | 0,0254 | 0,0228 | 0,024 | 0,0175 | | | 0,01859 | | |
| **2_Or_L** | 0,01685 | 0,01548 | 0,0144 | 0,016 | 0,017 | 0,0138 | 0,0147 | 0,014 | 0,0133 | 0,0129 | 0,0157 | 0,015 | 0,0152 | | 0,0146 | 0,0133 | | 0,0164 |
| **2_Or_R** | 0,0168 | 0,01533 | 0,0129 | 0,016 | 0,017 | | 0,0132 | 0,016 | 0,0154 | 0,0142 | 0,0139 | 0,015 | 0,0163 | | 0,0157 | 0,01487 | 0,0138 | |
| **2_AM_L** | 0,00472 | 0,00851 | 0,0092 | 0,007 | 0,007 | 0,0075 | 0,0067 | | 0,0066 | 0,003 | | 0,004 | | 0,0073 | 0,0064 | 0,00473 | 0,0039 | 0,0055 |
| **2_AM_R** | 0,00221 | 0,00797 | 0,0075 | 0,005 | 0,005 | 0,0053 | 0,0047 | 0,006 | | 0,0025 | 0,0028 | 0,004 | | | 0,0051 | 0,00477 | 0,0053 | 0,0053 |
| **2_Pir_L** | 0,00138 | 0,00179 | 0,0019 | 0,002 | 0,002 | 0,002 | 0,0018 | 0,002 | 0,0015 | 0,002 | 0,0025 | 0,002 | | 0,0012 | 0,0013 | 0,00145 | 0,0011 | 0,0013 |
| **2_Pir_R** | 0,00175 | 0,00183 | 0,0013 | 0,001 | 0,002 | 0,002 | 0,0019 | 0,001 | 0,001 | 0,0016 | 0,0021 | 0,002 | 0,001 | 0,0011 | 0,0011 | 0,00112 | 0,001 | 0,0009 |
| **2_M1_L** | 0,0882 | 0,09605 | 0,081 | 0,083 | 0,094 | 0,0882 | 0,0933 | 0,083 | 0,0806 | 0,0808 | 0,0804 | 0,08 | 0,1017 | 0,1069 | 0,1039 | 0,09412 | 0,0908 | 0,0875 |
| **2_M1_R** | 0,09557 | 0,08516 | 0,0826 | 0,085 | 0,096 | 0,0987 | 0,0921 | 0,089 | 0,0793 | 0,0838 | 0,0894 | 0,079 | 0,0976 | 0,1063 | 0,1022 | 0,08436 | 0,0852 | 0,0951 |
| **2_S2_L** | 0,08601 | 0,09877 | 0,0904 | 0,091 | 0,097 | 0,0951 | 0,0964 | 0,094 | 0,082 | 0,094 | 0,0885 | 0,076 | | 0,0995 | 0,0962 | 0,08856 | 0,0904 | 0,0842 |
| **2_S2_R** | 0,08748 | 0,09659 | 0,0955 | 0,081 | 0,094 | 0,0964 | 0,0906 | 0,092 | 0,093 | 0,086 | 0,0817 | 0,085 | 0,0929 | 0,0984 | 0,0984 | 0,08856 | 0,0918 | 0,0809 |
| **2_mt_L** | 0,00024 | 0,0003 | 0,0002 | 2E-04 | 3E-04 | 0,0002 | 0,0003 | 3E-04 | 0,0003 | 0,0002 | | 2E-04 | 0,0002 | 0,0003 | 0,0003 | 0,00023 | 0,0002 | 0,0002 |
| **2_mt_R** | 0,00019 | 0,0003 | 0,0002 | 2E-04 | 3E-04 | 0,0003 | 0,0003 | 3E-04 | 0,0003 | 0,0002 | | 2E-04 | 0,0002 | 0,0003 | 0,0003 | 0,00021 | 0,0002 | 0,0002 |
| **2_ic_L** | 0,00906 | 0,01051 | 0,0117 | 0,008 | 0,01 | 0,0107 | 0,0099 | 0,01 | 0,0098 | 0,0073 | 0,0075 | 0,006 | | 0,0087 | 0,0092 | 0,00742 | 0,0074 | 0,0074 |
| **2_ic_R** | 0,00802 | 0,01004 | 0,011 | 0,008 | 0,009 | 0,0099 | 0,0104 | 0,009 | 0,0095 | 0,0058 | 0,0053 | 0,008 | 0,0095 | 0,0086 | 0,0089 | 0,00697 | 0,0069 | 0,007 |
| **2_opt_L** | 0,00124 | 0,00143 | 0,0014 | 0,001 | 0,002 | 0,0027 | 0,0013 | 0,001 | 0,0016 | | | | 0,0013 | 0,0012 | 0,0014 | | | |
| **2_opt_R** | 0,00084 | 0,0011 | 0,0016 | 0,001 | 0,001 | 0,0012 | 0,0011 | 0,002 | 0,0012 | | | | 0,0015 | 0,0012 | 0,001 | | | |
| **2_fi_L** | 0,00404 | 0,00443 | 0,0038 | 0,003 | | 0,0041 | 0,0042 | 0,004 | 0,0036 | 0,0034 | 0,0035 | 0,003 | 0,0035 | 0,0035 | 0,0035 | 0,00314 | 0,0032 | 0,0032 |
| **2_fi_R** | 0,00405 | 0,00469 | 0,0042 | 0,003 | | 0,0032 | 0,0039 | 0,004 | 0,0035 | 0,0029 | 0,0034 | 0,003 | 0,0036 | 0,0033 | 0,0037 | 0,00304 | | |
| **2_Hb_L** | 0,00069 | 0,00087 | 0,0009 | 1E-03 | 9E-04 | 0,0008 | 0,0008 | 9E-04 | 0,0009 | 0,0008 | 0,0004 | 6E-04 | 0,0006 | 0,0005 | 0,0005 | 0,00048 | 0,0005 | 0,0005 |
| **2_Hb_R** | 0,00062 | 0,00098 | 0,0009 | 0,001 | 9E-04 | 0,0007 | 0,0008 | 8E-04 | 0,0009 | 0,0008 | 0,0004 | 6E-04 | 0,0006 | 0,0005 | 0,0005 | 0,00046 | 0,0005 | 0,0005 |

**Supplementary Table 4** Raw hippocampal measurements for male and female KO mice

| Barcode | Mice | Sex | CA1_length_left | CA1_length_right | CA2 length left | CA2_length_right |
|---|---|---|---|---|---|---|
| 2_A00001388 | WT | Female | 0,134 | 0,153 | 0,017 | 0,011 |
| 2_A00001389 | WT | Female | 0,103 | 0,106 | 0,021 | 0,024 |
| 2_A00001390 | WT | Female | 0,135 | 0,114 | 0,026 | 0,024 |
| 2_A00001391 | WT | Male | 0,129 | 0,122 | 0,019 | 0,024 |
| 2_A00001392 | WT | Male | 0,166 | | 0,026 | |
| 2_A00001393 | WT | Male | 0,152 | 0,194 | 0,023 | 0,023 |
| 2_A00001394 | HOM | Female | 0,106 | 0,106 | 0,018 | 0,017 |
| 2_A00001395 | HOM | Female | 0,093 | 0,111 | 0,017 | 0,013 |
| 2_A00001396 | HOM | Female | 0,106 | 0,089 | 0,02 | 0,019 |
| 2_A00001397 | HOM | Male | 0,129 | | 0,02 | |
| 2_A00001398 | HOM | Male | 0,141 | 0,157 | 0,02 | 0,024 |
| 2_A00001399 | HOM | Male | 0,133 | 0,137 | 0,019 | 0,021 |

| Barcode | Mice | Sex | CA3_length_left | CA3_length_right | CA1_area_left | CA1_area_right |
|---|---|---|---|---|---|---|
| 2_A00001388 | WT | Female | 0,053 | 0,054 | 0,0007938 | 0,0008252 |
| 2_A00001389 | WT | Female | 0,048 | 0,037 | 0,000657 | 0,0006178 |
| 2_A00001390 | WT | Female | 0,08 | 0,061 | 0,0007705 | 0,0007302 |
| 2_A00001391 | WT | Male | 0,065 | 0,093 | 0,0006413 | 0,0005881 |
| 2_A00001392 | WT | Male | 0,088 | | 0,0009633 | |
| 2_A00001393 | WT | Male | 0,077 | 0,057 | 0,0006637 | 0,0007984 |
| 2_A00001394 | HOM | Female | 0,066 | 0,062 | 0,0004438 | 0,0004846 |
| 2_A00001395 | HOM | Female | 0,073 | 0,076 | 0,0004233 | 0,0004807 |
| 2_A00001396 | HOM | Female | 0,063 | 0,037 | 0,00043904 | 0,0004339 |
| 2_A00001397 | HOM | Male | 0,061 | | 0,0005799 | |
| 2_A00001398 | HOM | Male | 0,065 | 0,049 | 0,0006964 | 0,0007389 |
| 2_A00001399 | HOM | Male | 0,068 | 0,049 | 0,0006025 | 0,0006086 |

| Barcode | Mice | Sex | CA2_area_left | CA2_area_right | CA3_area_left | CA3_area_right |
|---|---|---|---|---|---|---|
| 2_A00001388 | WT | Female | 0,0001217 | 0,0001001 | 0,0005169 | 0,0004782 |
| 2_A00001389 | WT | Female | 0,0001759 | 0,0001993 | 0,0005439 | 0,0003946 |
| 2_A00001390 | WT | Female | 0,0001977 | 0,0001798 | 0,0008362 | 0,0006217 |
| 2_A00001391 | WT | Male | 0,0001218 | 0,0001725 | 0,000572 | 0,0006948 |
| 2_A00001392 | WT | Male | 0,0001774 | | 0,0009443 | |
| 2_A00001393 | WT | Male | 0,0001386 | 0,0001328 | 0,0005259 | 0,0004784 |
| 2_A00001394 | HOM | Female | 0,0001094 | 0,00009927 | 0,0007825 | 0,000525 |
| 2_A00001395 | HOM | Female | 0,0001166 | 0,00013934 | 0,0005244 | 0,000674 |
| 2_A00001396 | HOM | Female | 0,0001142 | 0,0001673 | 0,0004757 | 0,0003773 |
| 2_A00001397 | HOM | Male | 0,0001365 | | 0,0005939 | |
| 2_A00001398 | HOM | Male | 0,0001525 | 0,0001951 | 0,0004867 | 0,0003587 |
| 2_A00001399 | HOM | Male | 0,0001482 | 0,0001419 | 0,0005214 | 0,0003859 |

| Barcode | Mice | Sex | GrDG_length_up | GrDG_length_uppe | GrDG_length_lo | GrDG_length_lower_right |
|---|---|---|---|---|---|---|
| 2_A00001388 | WT | Female | 0,066 | 0,08 | 0,099 | 0,101 |
| 2_A00001389 | WT | Female | | | 0,112 | 0,111 |
| 2_A00001390 | WT | Female | 0,069 | 0,051 | 0,102 | 0,094 |
| 2_A00001391 | WT | Male | 0,064 | 0,069 | 0,086 | 0,086 |
| 2_A00001392 | WT | Male | 0,091 | | 0,106 | |
| 2_A00001393 | WT | Male | 0,076 | 0,094 | 0,093 | 0,092 |
| 2_A00001394 | HOM | Female | 0,057 | 0,059 | 0,083 | 0,079 |
| 2_A00001395 | HOM | Female | 0,047 | 0,06 | 0,09 | 0,095 |
| 2_A00001396 | HOM | Female | 0,06 | | 0,087 | |
| 2_A00001397 | HOM | Male | 0,059 | | 0,074 | |
| 2_A00001398 | HOM | Male | 0,06 | 0,053 | 0,071 | 0,074 |
| 2_A00001399 | HOM | Male | 0,067 | 0,069 | 0,075 | 0,071 |

| Barcode | Mice | Sex | GrDG_area_uppe | GrDG_area_upper | GrDG_area_low | GrDG_area_lower_right |
|---|---|---|---|---|---|---|
| 2_A00001388 | WT | Female | 2,52E-04 | 3,31E-04 | 5,61E-04 | 4,63E-04 |
| 2_A00001389 | WT | Female | | | 6,91E-04 | 6,96E-04 |
| 2_A00001390 | WT | Female | 3,67E-04 | 2,27E-04 | 6,81E-04 | 6,44E-04 |
| 2_A00001391 | WT | Male | 2,99E-04 | 3,23E-04 | 5,30E-04 | 5,60E-04 |
| 2_A00001392 | WT | Male | 5,17E-04 | | 6,54E-04 | |
| 2_A00001393 | WT | Male | 3,84E-04 | 5,14E-04 | 5,03E-04 | 5,35E-04 |
| 2_A00001394 | HOM | Female | 2,40E-04 | 2,38E-04 | 5,05E-04 | 5,02E-04 |
| 2_A00001395 | HOM | Female | 1,67E-04 | 2,55E-04 | 5,37E-04 | 5,90E-04 |
| 2_A00001396 | HOM | Female | 2,72E-04 | | 5,52E-04 | |
| 2_A00001397 | HOM | Male | 2,51E-04 | | 4,79E-04 | |
| 2_A00001398 | HOM | Male | 3,05E-04 | 2,82E-04 | 4,99E-04 | 4,45E-04 |
| 2_A00001399 | HOM | Male | 3,71E-04 | 3,69E-04 | 4,75E-04 | 4,39E-04 |

**Supplementary Table 5** - Table reporting the statistics of the 445 differentially expressed genes detected with DESeq2. Genes are ordered according to their adjusted P value.

| gene | baseMean | log2FoldChange | lfcSE | stat | pvalue | padj |
|---|---|---|---|---|---|---|
| Rnf213 | 1557,32101 | -2,365374232 | 0,18656155 | -12,834168 | 1,06E-37 | 1,79E-33 |
| Iigp1 | 179,268122 | -2,636324186 | 0,21247444 | -12,181671 | 3,89E-34 | 3,31E-30 |
| Parp14 | 183,327807 | -2,145664176 | 0,18133094 | -11,69365 | 1,37E-31 | 7,78E-28 |
| Irgm1 | 172,903575 | -2,118137894 | 0,18090294 | -11,541246 | 8,17E-31 | 3,47E-27 |
| Mx1 | 75,4009844 | -2,794964046 | 0,21839619 | -11,079921 | 1,57E-28 | 5,33E-25 |
| Oasl2 | 256,524942 | -2,248907955 | 0,21234974 | -10,84641 | 2,07E-27 | 5,03E-24 |
| Irf7 | 103,094898 | -2,238220056 | 0,19847629 | -10,857983 | 1,83E-27 | 5,03E-24 |
| Isg15 | 89,9108619 | -2,696793214 | 0,2204231 | -10,718381 | 8,35E-27 | 1,77E-23 |
| H2-D1 | 569,048211 | -1,894796892 | 0,18523215 | -10,277128 | 8,94E-25 | 1,69E-21 |
| Ifih1 | 149,031673 | -2,083698737 | 0,20956389 | -9,9593678 | 2,30E-23 | 3,90E-20 |
| Gbp7 | 119,004213 | -1,807687156 | 0,18291605 | -9,7252742 | 2,35E-22 | 3,63E-19 |
| Rsad2 | 80,5739543 | -2,230197757 | 0,22528665 | -9,6565109 | 4,61E-22 | 6,53E-19 |
| Lgals3bp | 290,894613 | -1,888471871 | 0,19742914 | -9,611017 | 7,18E-22 | 9,39E-19 |
| Oas2 | 69,649064 | -2,11278806 | 0,21470124 | -9,4435429 | 3,60E-21 | 4,37E-18 |
| Bst2 | 83,294053 | -2,082374759 | 0,21751602 | -9,4327308 | 4,00E-21 | 4,52E-18 |
| Stat1 | 249,23963 | -1,702207496 | 0,18167946 | -9,3859577 | 6,23E-21 | 6,62E-18 |
| Cxcl10 | 53,9832749 | -2,439377309 | 0,22638913 | -9,2596383 | 2,05E-20 | 2,05E-17 |
| Ifit2 | 102,615448 | -1,749325193 | 0,18850721 | -9,1320311 | 6,72E-20 | 6,34E-17 |
| Xaf1 | 96,2530475 | -1,578514778 | 0,17368455 | -8,9639699 | 3,13E-19 | 2,80E-16 |
| Zc3hav1 | 194,722752 | -1,751810422 | 0,19654911 | -8,9100928 | 5,10E-19 | 4,15E-16 |
| Mx2 | 87,248637 | -1,938960293 | 0,21688885 | -8,9093379 | 5,13E-19 | 4,15E-16 |
| H2-K1 | 321,765772 | -1,706344403 | 0,19466663 | -8,8138875 | 1,21E-18 | 9,33E-16 |
| Trim25 | 161,023683 | -1,645518441 | 0,18628227 | -8,806569 | 1,29E-18 | 9,53E-16 |
| Ifit3 | 109,571425 | -1,880525265 | 0,21500306 | -8,745046 | 2,23E-18 | 1,58E-15 |
| H2-Q4 | 141,968077 | -1,833060538 | 0,22356748 | -8,5310105 | 1,45E-17 | 9,86E-15 |
| Cdkn1a | 185,124617 | -1,80236693 | 0,21684832 | -8,5170977 | 1,64E-17 | 1,07E-14 |
| H2-T23 | 134,11801 | -1,796017034 | 0,21355176 | -8,5107841 | 1,73E-17 | 1,09E-14 |
| Slfn5 | 92,183479 | -1,75349044 | 0,20279119 | -8,4812404 | 2,23E-17 | 1,35E-14 |
| Gm4951 | 47,8178154 | -2,090901537 | 0,22358105 | -8,4232425 | 3,66E-17 | 2,15E-14 |
| Usp18 | 71,7438795 | -1,821767827 | 0,22415036 | -8,2414505 | 1,70E-16 | 9,63E-14 |
| Ddx60 | 161,916176 | -1,736404364 | 0,22189963 | -8,1283015 | 4,35E-16 | 2,39E-13 |
| Tap1 | 123,954758 | -1,669821641 | 0,20683509 | -8,1132343 | 4,93E-16 | 2,62E-13 |
| Ifit1 | 241,470551 | -1,448857732 | 0,22568354 | -7,8936453 | 2,93E-15 | 1,51E-12 |
| Gm20559 | 58,3471264 | -1,778497148 | 0,22269947 | -7,8401249 | 4,50E-15 | 2,25E-12 |
| Stat2 | 425,52988 | -1,231094111 | 0,15844458 | -7,7752496 | 7,53E-15 | 3,65E-12 |
| Gbp3 | 108,844472 | -1,601889262 | 0,21187227 | -7,6318446 | 2,31E-14 | 1,09E-11 |
| Samd9l | 129,499639 | -1,656046284 | 0,2237901 | -7,6166157 | 2,60E-14 | 1,20E-11 |
| Trim30a | 84,2988381 | -1,563255189 | 0,20322559 | -7,5975004 | 3,02E-14 | 1,35E-11 |
| Eif2ak2 | 136,012356 | -1,578531775 | 0,20911204 | -7,5777148 | 3,52E-14 | 1,53E-11 |
| Slfn8 | 55,498026 | -1,590230175 | 0,20909571 | -7,4928404 | 6,74E-14 | 2,86E-11 |
| Oas1b | 40,7589393 | -1,695493083 | 0,22077072 | -7,4459676 | 9,62E-14 | 3,99E-11 |
| Ifi213 | 33,3181103 | -1,776275826 | 0,22596544 | -7,3489274 | 2,00E-13 | 8,08E-11 |

| | | | | | | |
|---|---|---|---|---|---|---|
| Stac2 | 411,371997 | -1,313225089 | 0,18030234 | -7,2920196 | 3,05E-13 | 1,20E-10 |
| Tapbp | 443,242405 | -1,540223707 | 0,21189739 | -7,2891043 | 3,12E-13 | 1,20E-10 |
| Robo3 | 615,706866 | -1,421284654 | 0,19710546 | -7,2626768 | 3,80E-13 | 1,43E-10 |
| Ifi44 | 40,3584411 | -1,68048971 | 0,22620998 | -7,218149 | 5,27E-13 | 1,95E-10 |
| Ddx58 | 141,369908 | -1,550619357 | 0,22023533 | -7,1334969 | 9,79E-13 | 3,54E-10 |
| AW011738 | 92,3452611 | -1,36998986 | 0,19141029 | -7,0830578 | 1,41E-12 | 4,99E-10 |
| Igtp | 86,1677125 | -1,517175374 | 0,21621453 | -7,0791057 | 1,45E-12 | 5,03E-10 |
| Parp9 | 82,3241386 | -1,42948728 | 0,20035047 | -7,0664905 | 1,59E-12 | 5,40E-10 |
| H2-T22 | 151,586163 | -1,232202627 | 0,17499747 | -7,0337745 | 2,01E-12 | 6,70E-10 |
| Parp12 | 142,197885 | -1,086604686 | 0,15406812 | -7,0263886 | 2,12E-12 | 6,92E-10 |
| Gbp5 | 49,3891726 | -1,585396069 | 0,22062585 | -7,0227864 | 2,17E-12 | 6,97E-10 |
| F830016B08 | 41,6474769 | -1,601208559 | 0,22693773 | -7,0011974 | 2,54E-12 | 7,98E-10 |
| Rnf10 | 879,063166 | -1,03715848 | 0,15127611 | -6,852201 | 7,27E-12 | 2,25E-09 |
| Dtx3l | 143,371855 | -1,256334189 | 0,18517501 | -6,7611726 | 1,37E-11 | 4,15E-09 |
| Znfx1 | 485,021375 | -0,808904962 | 0,11976041 | -6,7541539 | 1,44E-11 | 4,21E-09 |
| Ifit3b | 64,1942985 | -1,477153457 | 0,21908168 | -6,7564929 | 1,41E-11 | 4,21E-09 |
| Trim34a | 45,0374163 | -1,425353534 | 0,21322355 | -6,5960266 | 4,22E-11 | 1,22E-08 |
| Gm6548 | 37,4945903 | -1,428627354 | 0,21322505 | -6,5362611 | 6,31E-11 | 1,79E-08 |
| Helz2 | 141,790345 | -1,274933339 | 0,19664839 | -6,507484 | 7,64E-11 | 2,13E-08 |
| Oas3 | 24,3518046 | -1,588642021 | 0,22843234 | -6,4670027 | 1,00E-10 | 2,74E-08 |
| Ifi204 | 22,9778634 | -1,583631084 | 0,22845245 | -6,4630195 | 1,03E-10 | 2,77E-08 |
| C1qa | 274,894264 | -1,256276818 | 0,19452648 | -6,4520219 | 1,10E-10 | 2,93E-08 |
| Cd274 | 37,3130229 | -1,437235432 | 0,22077831 | -6,3891805 | 1,67E-10 | 4,36E-08 |
| C4b | 1419,56325 | -1,087626324 | 0,17248697 | -6,3320037 | 2,42E-10 | 6,23E-08 |
| Lag3 | 85,6726347 | -1,39842368 | 0,22035202 | -6,2986124 | 3,00E-10 | 7,61E-08 |
| Pik3ap1 | 64,859957 | -1,273908358 | 0,19946451 | -6,2931432 | 3,11E-10 | 7,77E-08 |
| Gm19410 | 360,573561 | -0,905068836 | 0,14375405 | -6,2839062 | 3,30E-10 | 8,13E-08 |
| Uba7 | 119,173784 | -1,309064919 | 0,21295107 | -6,272725 | 3,55E-10 | 8,61E-08 |
| Psmb8 | 42,2972965 | -1,382161558 | 0,22553718 | -6,1872571 | 6,12E-10 | 1,46E-07 |
| C1qc | 288,109386 | -1,25151414 | 0,20151116 | -6,1755136 | 6,59E-10 | 1,56E-07 |
| Herc6 | 82,4335943 | -1,303051861 | 0,21193938 | -6,1539076 | 7,56E-10 | 1,76E-07 |
| Trim21 | 37,5625321 | -1,392620253 | 0,22868827 | -6,151026 | 7,70E-10 | 1,77E-07 |
| Phf11d | 20,0641 | -1,500174473 | 0,22773151 | -6,0922118 | 1,11E-09 | 2,52E-07 |
| Ube2l6 | 39,3255428 | -1,380425957 | 0,22630467 | -6,00724 | 1,89E-09 | 4,22E-07 |
| Ptprc | 104,040234 | -1,236312522 | 0,20783039 | -5,9523346 | 2,64E-09 | 5,83E-07 |
| Car12 | 250,669114 | -0,994933121 | 0,16767266 | -5,9211683 | 3,20E-09 | 6,96E-07 |
| Trim30d | 43,2415161 | -1,315060175 | 0,22226773 | -5,8828753 | 4,03E-09 | 8,67E-07 |
| Prkca | 2494,96531 | 0,7150363 | 0,1219593 | 5,86593709 | 4,47E-09 | 9,48E-07 |
| Ifi35 | 34,263546 | -1,319919458 | 0,22670623 | -5,8245274 | 5,73E-09 | 1,20E-06 |
| Parp10 | 50,6650101 | -1,270029054 | 0,2216474 | -5,8143417 | 6,09E-09 | 1,26E-06 |
| Hrh3 | 176,392565 | -1,149929041 | 0,19725876 | -5,8091073 | 6,28E-09 | 1,29E-06 |
| Irf9 | 156,64812 | -1,0444075 | 0,18133894 | -5,7708295 | 7,89E-09 | 1,60E-06 |
| Rasgrf2 | 197,078137 | 1,026398237 | 0,17772022 | 5,76839939 | 8,00E-09 | 1,60E-06 |
| Axl | 224,969848 | -1,019448546 | 0,1765767 | -5,7621096 | 8,31E-09 | 1,64E-06 |
| Oasl1 | 17,8529501 | -1,394655338 | 0,2276374 | -5,7302422 | 1,00E-08 | 1,96E-06 |
| Rtp4 | 36,5331411 | -1,283506454 | 0,22790206 | -5,6684191 | 1,44E-08 | 2,78E-06 |

| | | | | | | |
|---|---|---|---|---|---|---|
| C1qb | 223,838632 | -1,094989474 | 0,19421971 | -5,6282225 | 1,82E-08 | 3,48E-06 |
| Il17ra | 138,554081 | -0,940871102 | 0,16736681 | -5,6140477 | 1,98E-08 | 3,73E-06 |
| Camta1 | 1844,94547 | 0,583847211 | 0,10430272 | 5,59713357 | 2,18E-08 | 4,07E-06 |
| Ifi203 | 32,9970132 | -1,245476247 | 0,22266325 | -5,5598772 | 2,70E-08 | 4,98E-06 |
| Siglec1 | 18,0106759 | -1,298354665 | 0,22703551 | -5,5564406 | 2,75E-08 | 4,98E-06 |
| Ap1ar | 344,029879 | 0,761207858 | 0,13682726 | 5,5575022 | 2,74E-08 | 4,98E-06 |
| Plxnd1 | 314,071127 | -0,805302567 | 0,14558354 | -5,5265043 | 3,27E-08 | 5,84E-06 |
| Plekha4 | 29,0498035 | -1,25943844 | 0,22622066 | -5,5213044 | 3,36E-08 | 5,89E-06 |
| H2-T24 | 31,1682895 | -1,242017772 | 0,22188892 | -5,5218336 | 3,35E-08 | 5,89E-06 |
| Slfn2 | 21,9361724 | -1,268054996 | 0,22859203 | -5,4214419 | 5,91E-08 | 1,02E-05 |
| Ctsz | 152,699623 | -0,985694083 | 0,18234661 | -5,4043128 | 6,51E-08 | 1,12E-05 |
| Gm31614 | 19,6247017 | -1,209649504 | 0,22487254 | -5,3965815 | 6,79E-08 | 1,15E-05 |
| Mras | 773,788523 | 0,640260618 | 0,11888404 | 5,38333698 | 7,31E-08 | 1,23E-05 |
| Nlrc5 | 220,177801 | -1,111015194 | 0,22912528 | -5,3796707 | 7,46E-08 | 1,24E-05 |
| Oas1a | 22,6330603 | -1,248565227 | 0,22881787 | -5,3777754 | 7,54E-08 | 1,24E-05 |
| Itgax | 41,0609855 | -1,217727743 | 0,22737082 | -5,3406794 | 9,26E-08 | 1,48E-05 |
| Dhx58 | 28,1151086 | -1,209117674 | 0,22317787 | -5,3444198 | 9,07E-08 | 1,48E-05 |
| Ifi27l2a | 27,1039277 | -1,193974116 | 0,22888758 | -5,3394277 | 9,32E-08 | 1,48E-05 |
| Golm1 | 283,247206 | 0,854889158 | 0,16026266 | 5,34028464 | 9,28E-08 | 1,48E-05 |
| Gbp2 | 55,8857147 | -1,176198074 | 0,22593704 | -5,3115028 | 1,09E-07 | 1,71E-05 |
| Tap2 | 94,1492421 | -1,136225681 | 0,21585905 | -5,3093333 | 1,10E-07 | 1,71E-05 |
| Ctss | 330,719499 | -1,103850037 | 0,20633306 | -5,2909653 | 1,22E-07 | 1,88E-05 |
| Gbp4 | 48,5151892 | -1,133280093 | 0,22782271 | -5,2723013 | 1,35E-07 | 2,06E-05 |
| Ifi209 | 15,1480354 | -1,278529048 | 0,22851058 | -5,2204985 | 1,78E-07 | 2,67E-05 |
| Gm4841 | 15,9179361 | -1,508513065 | 0,22482608 | -5,2199152 | 1,79E-07 | 2,67E-05 |
| Zbp1 | 26,3432383 | -1,102750068 | 0,2283932 | -5,2017049 | 1,97E-07 | 2,92E-05 |
| Camk1d | 1321,46437 | 0,594947365 | 0,11496592 | 5,17565455 | 2,27E-07 | 3,33E-05 |
| Nptx2 | 292,468878 | -1,041097412 | 0,2036764 | -5,1727079 | 2,31E-07 | 3,35E-05 |
| Shisa5 | 212,251818 | -0,804932771 | 0,15636502 | -5,1473696 | 2,64E-07 | 3,80E-05 |
| Cd22 | 15,0760089 | -1,234075031 | 0,22749669 | -5,1282903 | 2,92E-07 | 4,17E-05 |
| Oas1g | 14,0544001 | -1,336684698 | 0,22484801 | -5,1151783 | 3,13E-07 | 4,44E-05 |
| Etv5 | 540,565009 | -0,597632017 | 0,1177045 | -5,0775493 | 3,82E-07 | 5,37E-05 |
| Sp100 | 42,04132 | -1,067986802 | 0,20828098 | -5,0757731 | 3,86E-07 | 5,37E-05 |
| Apobec3 | 58,5387278 | -1,007635684 | 0,19896126 | -5,0707104 | 3,96E-07 | 5,47E-05 |
| Stum | 3156,55406 | 0,71786724 | 0,14196137 | 5,05803822 | 4,24E-07 | 5,80E-05 |
| Ifitm3 | 41,5800986 | -1,143268778 | 0,22694478 | -5,0487311 | 4,45E-07 | 6,04E-05 |
| Tor3a | 58,8362947 | -1,110941731 | 0,21857869 | -5,0467694 | 4,49E-07 | 6,06E-05 |
| Pml | 105,945967 | -0,828487252 | 0,16477581 | -5,0242155 | 5,05E-07 | 6,76E-05 |
| Phlda3 | 95,9160999 | -1,05745444 | 0,21140234 | -5,0003463 | 5,72E-07 | 7,59E-05 |
| Ltbp1 | 57,1960362 | -0,982215108 | 0,19453581 | -4,9978498 | 5,80E-07 | 7,63E-05 |
| Tcirg1 | 163,928142 | -1,003059355 | 0,20257812 | -4,9873083 | 6,12E-07 | 8,00E-05 |
| Nos1 | 491,481032 | 0,940569172 | 0,18915423 | 4,97089933 | 6,66E-07 | 8,64E-05 |
| Irf8 | 76,6995245 | -1,003646055 | 0,20227913 | -4,9437359 | 7,66E-07 | 9,86E-05 |
| Bag3 | 66,948684 | -1,066419782 | 0,21458597 | -4,9298938 | 8,23E-07 | 0,00010508 |
| Gbp9 | 55,7098779 | -1,049685563 | 0,21406469 | -4,9239005 | 8,48E-07 | 0,00010755 |
| Grn | 236,814881 | -0,770473987 | 0,15703137 | -4,8999775 | 9,58E-07 | 0,0001206 |

| | | | | | | |
|---|---|---|---|---|---|---|
| Aspg | 33,0445628 | -1,086210624 | 0,22741663 | -4,8534679 | 1,21E-06 | 0,00015154 |
| Crlf1 | 18,7335297 | 1,027431382 | 0,22453708 | 4,83841057 | 1,31E-06 | 0,00016228 |
| Vim | 149,413075 | -0,966357983 | 0,19928279 | -4,8325899 | 1,35E-06 | 0,00016589 |
| Ecel1 | 41,67843 | -1,029638253 | 0,2288276 | -4,8108587 | 1,50E-06 | 0,00018366 |
| Pcdh20 | 780,19173 | 0,618008029 | 0,1291541 | 4,78626785 | 1,70E-06 | 0,0002047 |
| Hcn3 | 89,6880812 | -0,820149777 | 0,17146938 | -4,7712805 | 1,83E-06 | 0,00021746 |
| Trim12a | 59,2641055 | -0,97528074 | 0,20424092 | -4,7667904 | 1,87E-06 | 0,00022081 |
| Ak5 | 331,125331 | 0,631489371 | 0,1326753 | 4,75973481 | 1,94E-06 | 0,0002271 |
| Cd44 | 42,3468868 | -1,087150423 | 0,22784691 | -4,7560773 | 1,97E-06 | 0,00022966 |
| Ccl5 | 12,765379 | -1,168337346 | 0,22328504 | -4,7425286 | 2,11E-06 | 0,0002439 |
| 9330162G02 | 1163,00215 | 0,981969139 | 0,20723678 | 4,69914691 | 2,61E-06 | 0,00029986 |
| Ctsd | 1074,24714 | -0,858802036 | 0,18331582 | -4,6851519 | 2,80E-06 | 0,00031894 |
| Ogfr | 180,354313 | -0,816952574 | 0,17470224 | -4,6748816 | 2,94E-06 | 0,00033168 |
| Mmp2 | 29,0285324 | -1,051012123 | 0,2243197 | -4,6730297 | 2,97E-06 | 0,00033168 |
| Irf1 | 43,7993189 | -1,038190126 | 0,22614993 | -4,6742343 | 2,95E-06 | 0,00033168 |
| Chrna4 | 159,334966 | -0,793329813 | 0,17046348 | -4,6577934 | 3,20E-06 | 0,00035486 |
| Gfap | 2134,73992 | -0,902273995 | 0,1948139 | -4,6274662 | 3,70E-06 | 0,00040568 |
| Icam1 | 29,0005763 | -1,076667362 | 0,22896324 | -4,6231451 | 3,78E-06 | 0,00041157 |
| Nxph1 | 138,137538 | 0,941780119 | 0,20218506 | 4,61787332 | 3,88E-06 | 0,00041947 |
| Lratd2 | 124,371649 | -0,90097374 | 0,19503309 | -4,5781592 | 4,69E-06 | 0,00050433 |
| Arhgef6 | 114,871271 | 0,748631929 | 0,16331926 | 4,57527667 | 4,76E-06 | 0,0005081 |
| Tspo | 15,404595 | -1,058499763 | 0,22730019 | -4,5715019 | 4,84E-06 | 0,00051092 |
| Slc9a6 | 599,462864 | 0,771094269 | 0,16852836 | 4,57190117 | 4,83E-06 | 0,00051092 |
| Irgm2 | 73,3709689 | -1,000908617 | 0,22765285 | -4,5510308 | 5,34E-06 | 0,00055977 |
| Lgals9 | 40,3781055 | -1,007708776 | 0,22131753 | -4,547543 | 5,43E-06 | 0,00056563 |
| Cdc42bpg | 69,6177562 | -1,000207351 | 0,22042049 | -4,537954 | 5,68E-06 | 0,00058479 |
| H2-Q7 | 18,5534188 | -0,962118606 | 0,22626229 | -4,5251398 | 6,04E-06 | 0,00061763 |
| Pgm2l1 | 2470,23178 | 0,736102455 | 0,16282873 | 4,51906015 | 6,21E-06 | 0,00063182 |
| Fbln2 | 89,9900946 | -0,940922742 | 0,2081175 | -4,5087287 | 6,52E-06 | 0,00065553 |
| Laptm5 | 154,188832 | -0,761792181 | 0,16875639 | -4,506954 | 6,58E-06 | 0,00065715 |
| Gvin3 | 21,5204642 | -0,989209589 | 0,22884746 | -4,5011193 | 6,76E-06 | 0,00067054 |
| Cd300lf | 16,7843418 | -0,92885553 | 0,2240838 | -4,5001829 | 6,79E-06 | 0,00067054 |
| Slc24a3 | 1258,26803 | 0,79272904 | 0,17668163 | 4,49499983 | 6,96E-06 | 0,00068311 |
| Cmpk2 | 98,6241739 | -0,850867706 | 0,18892578 | -4,4855126 | 7,27E-06 | 0,00071012 |
| Tlr2 | 17,9248782 | -1,030459013 | 0,22797126 | -4,4719401 | 7,75E-06 | 0,00074813 |
| Slfn9 | 15,7256713 | -1,023063569 | 0,22833118 | -4,4727037 | 7,72E-06 | 0,00074813 |
| Nell2 | 3602,69384 | 0,61474952 | 0,13770144 | 4,4675101 | 7,91E-06 | 0,00075948 |
| Sp110 | 15,1012638 | -1,009619734 | 0,22842633 | -4,4620417 | 8,12E-06 | 0,00077474 |
| Hck | 19,4817175 | -1,011715738 | 0,2270611 | -4,4449522 | 8,79E-06 | 0,00082151 |
| Rnf114 | 267,649036 | -0,683630006 | 0,15367614 | -4,4463073 | 8,74E-06 | 0,00082151 |
| Vcam1 | 109,36495 | -0,980415018 | 0,21792172 | -4,4446935 | 8,80E-06 | 0,00082151 |
| Runx1 | 36,4438286 | -0,963410746 | 0,21502303 | -4,4322282 | 9,33E-06 | 0,00086573 |
| H2-Q5 | 18,9609844 | -0,94323252 | 0,22406089 | -4,415334 | 1,01E-05 | 0,00093109 |
| Tomm20 | 1086,7757 | 0,584976098 | 0,1328961 | 4,40063326 | 1,08E-05 | 0,00098048 |
| Ifi47 | 16,833364 | -0,956963417 | 0,2267705 | -4,39376 | 1,11E-05 | 0,00100663 |
| Fndc1 | 34,5367728 | 0,954162555 | 0,21916695 | 4,36997041 | 1,24E-05 | 0,00111686 |

| | | | | | | |
|---|---|---|---|---|---|---|
| Trim56 | 119,214151 | -0,915226734 | 0,20964784 | -4,3628318 | 1,28E-05 | 0,00114787 |
| Msn | 118,48596 | -0,888208225 | 0,2033532 | -4,3410088 | 1,42E-05 | 0,0012614 |
| Cyba | 23,612224 | -0,996706543 | 0,22545499 | -4,3372334 | 1,44E-05 | 0,00127657 |
| Isg20 | 9,91751492 | -0,961124298 | 0,2186433 | -4,2956806 | 1,74E-05 | 0,00151329 |
| Jcad | 885,899771 | -0,647321552 | 0,1507229 | -4,2938244 | 1,76E-05 | 0,00151436 |
| Npc2 | 195,227522 | -0,697065947 | 0,1624393 | -4,2885649 | 1,80E-05 | 0,00154283 |
| Psmb9 | 24,9931289 | -0,937975622 | 0,22892508 | -4,2742054 | 1,92E-05 | 0,00163741 |
| Dlx6os1 | 226,277197 | 0,625536847 | 0,14655009 | 4,26180375 | 2,03E-05 | 0,00172234 |
| Trim14 | 15,5596017 | -0,919989384 | 0,22562258 | -4,2538517 | 2,10E-05 | 0,00177581 |
| Hpcal4 | 1777,00855 | 0,619790476 | 0,14603319 | 4,23978475 | 2,24E-05 | 0,00188147 |
| Igsf3 | 84,785838 | -0,74193397 | 0,17506193 | -4,2355546 | 2,28E-05 | 0,0019078 |
| 6330403K07l | 826,195823 | -0,615586259 | 0,14584459 | -4,219177 | 2,45E-05 | 0,00203178 |
| Ifi27 | 233,913719 | -0,736267835 | 0,17563107 | -4,2001541 | 2,67E-05 | 0,00218889 |
| Trem2 | 95,803827 | -0,938778597 | 0,2220281 | -4,1851873 | 2,85E-05 | 0,00232698 |
| Gm37437 | 48,3535578 | 0,815513967 | 0,19275327 | 4,17990005 | 2,92E-05 | 0,00237035 |
| Gm48408 | 36,3254624 | 0,943506204 | 0,21973962 | 4,1661042 | 3,10E-05 | 0,00249451 |
| Gm16758 | 17,7102243 | 0,977465195 | 0,22826146 | 4,16698191 | 3,09E-05 | 0,00249451 |
| Tyro3 | 1337,07173 | -0,624270531 | 0,14990036 | -4,1646076 | 3,12E-05 | 0,00249908 |
| Cdc34 | 172,991906 | -0,686771513 | 0,16542235 | -4,1520421 | 3,30E-05 | 0,00262797 |
| Rnf39 | 27,4875817 | -0,956006392 | 0,22636756 | -4,1496521 | 3,33E-05 | 0,00264315 |
| Gpr12 | 102,19093 | 0,757817781 | 0,18258245 | 4,14686703 | 3,37E-05 | 0,00266305 |
| Edem1 | 275,996518 | -0,628714198 | 0,15166371 | -4,1438344 | 3,42E-05 | 0,00268604 |
| Dbp | 181,537498 | -0,782409025 | 0,18910904 | -4,1269492 | 3,68E-05 | 0,00287767 |
| Marcksl1 | 145,14461 | -0,846115598 | 0,20409882 | -4,1147412 | 3,88E-05 | 0,00300658 |
| Gm43196 | 9,25452959 | -0,938730243 | 0,21921301 | -4,1133685 | 3,90E-05 | 0,00301077 |
| Zfp92 | 21,2930916 | 0,924277427 | 0,22659695 | 4,09527999 | 4,22E-05 | 0,00322645 |
| Tent5c | 45,7004807 | -0,904493681 | 0,21980884 | -4,0886946 | 4,34E-05 | 0,00330452 |
| Rnf11 | 787,462469 | 0,637151007 | 0,15573723 | 4,08763044 | 4,36E-05 | 0,00330489 |
| Elk1 | 311,477189 | 0,694969718 | 0,1701871 | 4,07150073 | 4,67E-05 | 0,00351099 |
| Slc15a3 | 23,8821417 | -0,934269035 | 0,22810843 | -4,0511017 | 5,10E-05 | 0,00381012 |
| Smim43 | 27,1560691 | -0,926039965 | 0,22563812 | -4,048941 | 5,14E-05 | 0,00381651 |
| Ifi207 | 13,1396558 | -0,873015001 | 0,22280211 | -4,038129 | 5,39E-05 | 0,00397933 |
| St14 | 14,1456092 | -0,925813333 | 0,22759873 | -4,0324245 | 5,52E-05 | 0,00405955 |
| Col4a1 | 790,094451 | -0,834951821 | 0,20688199 | -4,030045 | 5,58E-05 | 0,00408319 |
| Adcy7 | 70,0267386 | -0,791712205 | 0,19726896 | -4,0274136 | 5,64E-05 | 0,0041059 |
| Plce1 | 129,985528 | -0,783768233 | 0,19523388 | -4,0232872 | 5,74E-05 | 0,00413437 |
| Flnc | 45,6550511 | -0,903755831 | 0,22629358 | -4,0169763 | 5,89E-05 | 0,00422522 |
| Cebpb | 64,2939176 | -0,881438571 | 0,21818875 | -4,0100646 | 6,07E-05 | 0,00433255 |
| Adamtsl3 | 25,8145237 | -0,907194045 | 0,22579635 | -4,0050076 | 6,20E-05 | 0,00440777 |
| Arl8a | 648,307892 | -0,583936562 | 0,14608055 | -3,9956955 | 6,45E-05 | 0,00456558 |
| H2-Q6 | 9,80926839 | -0,867137268 | 0,21887428 | -3,9842229 | 6,77E-05 | 0,00477196 |
| Fcgr1 | 22,5587115 | -0,911104298 | 0,22897779 | -3,9809579 | 6,86E-05 | 0,004818 |
| Ifi206 | 10,5675089 | -0,871038282 | 0,22249694 | -3,9746954 | 7,05E-05 | 0,004906 |
| Egr1 | 2233,86435 | -0,648307483 | 0,1635953 | -3,9748486 | 7,04E-05 | 0,004906 |
| Gm12250 | 10,8040441 | -0,760379959 | 0,21324487 | -3,9700239 | 7,19E-05 | 0,0049814 |
| Cxcl16 | 20,0982855 | -0,894678008 | 0,22894751 | -3,9691187 | 7,21E-05 | 0,0049814 |

| | | | | | | |
|---|---|---|---|---|---|---|
| Themis2 | 16,4197624 | -0,933633025 | 0,22774343 | -3,9680092 | 7,25E-05 | 0,00498281 |
| Rny1 | 208,874108 | -0,798322092 | 0,20016212 | -3,9661615 | 7,30E-05 | 0,00498281 |
| Cybb | 24,3296148 | -0,854781338 | 0,22768973 | -3,9664015 | 7,30E-05 | 0,00498281 |
| Gm43197 | 14,2073982 | -0,922562439 | 0,22854616 | -3,9571392 | 7,59E-05 | 0,00515404 |
| 9330175E14l | 9,19535023 | -0,863983258 | 0,21787012 | -3,9524371 | 7,74E-05 | 0,00523546 |
| Lrrk1 | 49,0687232 | -0,807831365 | 0,20592317 | -3,9405585 | 8,13E-05 | 0,0054798 |
| Dok1 | 23,6740638 | -0,900002129 | 0,22840279 | -3,9348086 | 8,33E-05 | 0,00559047 |
| Plek | 116,822497 | -0,773356948 | 0,19575929 | -3,9315254 | 8,44E-05 | 0,00564507 |
| Serping1 | 18,008353 | -0,867932021 | 0,22840232 | -3,9266006 | 8,62E-05 | 0,00573927 |
| Igfbp6 | 23,9621216 | -0,894298782 | 0,22881952 | -3,9180839 | 8,93E-05 | 0,00589956 |
| Egr3 | 1032,17099 | -0,654112863 | 0,16732932 | -3,9144532 | 9,06E-05 | 0,00596404 |
| Itgb5 | 236,570209 | -0,697951514 | 0,17835874 | -3,9135914 | 9,09E-05 | 0,00596404 |
| Ly86 | 42,2127 | -0,897686534 | 0,22748682 | -3,909033 | 9,27E-05 | 0,00605432 |
| Eid1 | 626,302963 | 0,733100285 | 0,18699677 | 3,9058955 | 9,39E-05 | 0,00610992 |
| Psmb10 | 60,4120486 | -0,839275697 | 0,216179 | -3,9026594 | 9,51E-05 | 0,00616858 |
| Cdk19 | 376,717868 | 0,613465385 | 0,15703286 | 3,90145431 | 9,56E-05 | 0,00617581 |
| Fcgr2b | 32,0644137 | -0,893824838 | 0,22667026 | -3,8967169 | 9,75E-05 | 0,00626876 |
| Htr1b | 295,006391 | -0,582601311 | 0,14948703 | -3,8957502 | 9,79E-05 | 0,00626876 |
| Platr25 | 75,7274984 | 0,763947623 | 0,19566834 | 3,89508883 | 9,82E-05 | 0,00626876 |
| Strip2 | 474,724189 | 0,753532919 | 0,19543585 | 3,89080978 | 9,99E-05 | 0,00635646 |
| Lgmn | 289,789548 | -0,627671966 | 0,16152612 | -3,8796427 | 0,00010461 | 0,00663063 |
| Calb1 | 283,035183 | 0,614983006 | 0,15838294 | 3,87465929 | 0,00010677 | 0,00669288 |
| Col18a1 | 24,8211775 | -0,898716591 | 0,22719618 | -3,8746619 | 0,00010677 | 0,00669288 |
| 9330121K16l | 187,611482 | 0,755483605 | 0,19450413 | 3,86468463 | 0,00011123 | 0,00694674 |
| Lacc1 | 34,4456334 | -0,829593797 | 0,21409825 | -3,8493324 | 0,00011844 | 0,00734286 |
| Dchs2 | 79,4466358 | 0,717229302 | 0,18802016 | 3,84820625 | 0,00011899 | 0,00734986 |
| Neurl3 | 13,031575 | -0,874324647 | 0,22738345 | -3,8435827 | 0,00012125 | 0,00745868 |
| B230303A05 | 88,927623 | 0,869973713 | 0,22510159 | 3,83375625 | 0,0001262 | 0,0076838 |
| Rasa4 | 41,9854319 | -0,855741086 | 0,22523383 | -3,8247147 | 0,00013092 | 0,00788652 |
| Gvin-ps7 | 53,7385908 | -0,831804074 | 0,21822568 | -3,8253135 | 0,00013061 | 0,00788652 |
| Zfp950 | 502,549259 | 0,588189972 | 0,15349914 | 3,82571297 | 0,00013039 | 0,00788652 |
| Creb3l1 | 33,5516107 | -0,852436352 | 0,22448123 | -3,8206712 | 0,00013309 | 0,00793257 |
| Mien1 | 120,381096 | -0,736281525 | 0,19234484 | -3,8223081 | 0,00013221 | 0,00793257 |
| Kcnq1ot1 | 2644,26247 | 0,588695656 | 0,15416336 | 3,81619735 | 0,00013552 | 0,00804947 |
| Kcnk6 | 29,4169685 | -0,844303184 | 0,22351021 | -3,8049069 | 0,00014186 | 0,00839629 |
| Perm1 | 9,37115233 | -0,825148403 | 0,22095162 | -3,7990646 | 0,00014524 | 0,00853719 |
| Itgal | 14,4503911 | -0,78721159 | 0,22316465 | -3,7929674 | 0,00014886 | 0,00871947 |
| Unc93b1 | 107,395443 | -0,732591396 | 0,19293964 | -3,7887147 | 0,00015143 | 0,00883959 |
| Cd72 | 7,64373846 | -0,810147049 | 0,20938662 | -3,7864985 | 0,00015279 | 0,00888822 |
| AU020206 | 73,7147868 | -0,794354136 | 0,21127752 | -3,7832284 | 0,00015481 | 0,00894459 |
| Slc9a4 | 60,5524962 | 0,813424712 | 0,21297183 | 3,77659025 | 0,00015899 | 0,0091242 |
| Gadd45g | 30,0891146 | -0,851212054 | 0,22838867 | -3,7713144 | 0,00016239 | 0,00928794 |
| Ucp2 | 81,0287633 | -0,750064671 | 0,20319296 | -3,7560837 | 0,00017259 | 0,00979144 |
| Gm19684 | 8,22753085 | -0,835234645 | 0,22096365 | -3,7556063 | 0,00017292 | 0,00979144 |
| Ftl1 | 440,824983 | -0,612544325 | 0,16326372 | -3,7503578 | 0,00017658 | 0,00993798 |
| Itpripl2 | 63,3588039 | -0,735293637 | 0,19461895 | -3,7414811 | 0,00018294 | 0,01018284 |

| | | | | | | |
|---|---|---|---|---|---|---|
| Igsf10 | 20,7858862 | -0,871023047 | 0,22875256 | -3,7341099 | 0,00018838 | 0,01036628 |
| Rhbdl3 | 182,084681 | -0,618497546 | 0,16578433 | -3,7335003 | 0,00018884 | 0,01036628 |
| Gm5970 | 10,1998693 | -0,774756657 | 0,2184652 | -3,7330476 | 0,00018918 | 0,01036628 |
| Rfk | 167,634358 | 0,619621095 | 0,16575197 | 3,73444473 | 0,00018813 | 0,01036628 |
| Gm44562 | 47,7363156 | 0,795305843 | 0,20898147 | 3,73140875 | 0,00019041 | 0,01040041 |
| Sort1 | 3615,8203 | 0,593988618 | 0,1595295 | 3,72529927 | 0,00019508 | 0,01059942 |
| Col5a1 | 73,2480228 | -0,801123869 | 0,2132861 | -3,7236247 | 0,00019638 | 0,01062406 |
| Col4a2 | 860,224534 | -0,828482578 | 0,22432785 | -3,7189286 | 0,00020007 | 0,01078915 |
| 1700037H04I | 124,937988 | -0,71658169 | 0,19246854 | -3,7168165 | 0,00020175 | 0,01082292 |
| Igfbp4 | 689,795526 | -0,721232856 | 0,19414466 | -3,7165394 | 0,00020197 | 0,01082292 |
| Piezo1 | 66,2822128 | -0,766095985 | 0,20881907 | -3,7064168 | 0,00021021 | 0,01122917 |
| Mapkapk3 | 21,5237886 | -0,840883665 | 0,22874982 | -3,7041315 | 0,00021212 | 0,01129535 |
| Szrd1 | 255,099719 | -0,62886378 | 0,16974777 | -3,7006849 | 0,00021502 | 0,01136033 |
| Mir22hg | 134,497643 | -0,614464712 | 0,16584753 | -3,7003024 | 0,00021534 | 0,01136033 |
| Tagln3 | 293,981372 | -0,680344162 | 0,18388216 | -3,7003078 | 0,00021534 | 0,01136033 |
| Slc9a5 | 51,4603705 | -0,752347975 | 0,2058415 | -3,6951558 | 0,00021975 | 0,01155705 |
| Fgl2 | 17,4359888 | -0,842803758 | 0,22873813 | -3,6847374 | 0,00022894 | 0,01200302 |
| Ppard | 154,017485 | -0,592047258 | 0,16074161 | -3,682892 | 0,0002306 | 0,01205308 |
| Kcns1 | 40,2474135 | -0,782184365 | 0,20998246 | -3,6811017 | 0,00023223 | 0,0120638 |
| Gm44559 | 70,5931589 | 0,748609462 | 0,20037484 | 3,67243743 | 0,00024025 | 0,01240454 |
| Trib1 | 125,969873 | -0,72826551 | 0,19866368 | -3,6715316 | 0,0002411 | 0,01241087 |
| Ano3 | 782,106677 | 0,64623602 | 0,17637003 | 3,66620287 | 0,00024618 | 0,01259589 |
| Eda2r | 18,8441564 | -0,83933767 | 0,22876324 | -3,6647033 | 0,00024763 | 0,01263187 |
| Wdfy4 | 51,4257581 | -0,725273235 | 0,19793015 | -3,6578714 | 0,00025432 | 0,01293446 |
| Nr2c2ap | 62,4071164 | -0,715938772 | 0,19578403 | -3,6560307 | 0,00025615 | 0,01298875 |
| Gm1043 | 220,761684 | 0,759403939 | 0,20825105 | 3,64917757 | 0,00026308 | 0,01330048 |
| Socs1 | 14,7020268 | -0,802523836 | 0,22105005 | -3,6481277 | 0,00026416 | 0,0133153 |
| Gap43 | 278,494193 | -0,656321989 | 0,17955444 | -3,6435912 | 0,00026886 | 0,01351222 |
| Itga5 | 22,8611979 | -0,818958576 | 0,22297598 | -3,6377327 | 0,00027505 | 0,01378246 |
| Lmo1 | 155,860591 | 0,627204483 | 0,1726107 | 3,62543381 | 0,00028848 | 0,01432852 |
| Gm42894 | 12,646929 | 0,835194287 | 0,22525926 | 3,61755885 | 0,00029739 | 0,01472841 |
| Ifi208 | 5,95395842 | -0,796029606 | 0,20511087 | -3,6141618 | 0,00030132 | 0,01483634 |
| A230103L15I | 259,625392 | 0,747096874 | 0,20519046 | 3,61275722 | 0,00030296 | 0,01484235 |
| mt-Nd2 | 3950,05612 | 0,630551326 | 0,17540026 | 3,61255874 | 0,00030319 | 0,01484235 |
| Slc5a5 | 62,5699947 | -0,754409166 | 0,20939571 | -3,6109288 | 0,0003051 | 0,01489303 |
| Adarb2 | 160,715786 | 0,658153531 | 0,18228581 | 3,5986393 | 0,00031989 | 0,01552546 |
| Anxa5 | 78,2725447 | -0,677833127 | 0,18827038 | -3,5871703 | 0,00033429 | 0,01613215 |
| Jak3 | 62,3719143 | -0,763947635 | 0,21448735 | -3,5763984 | 0,00034836 | 0,01676375 |
| B430305J03I | 38,4827186 | -0,798451476 | 0,22219965 | -3,5756125 | 0,00034941 | 0,01676669 |
| Lsp1 | 11,7446917 | -0,785538711 | 0,22526865 | -3,5741604 | 0,00035135 | 0,01680387 |
| Ccl12 | 5,95513943 | -0,815017828 | 0,20728243 | -3,5735588 | 0,00035216 | 0,01680387 |
| Gm45441 | 29,4473333 | -0,802345534 | 0,22159987 | -3,5720513 | 0,0003542 | 0,0168474 |
| Egr2 | 90,8725898 | -0,78502519 | 0,22764041 | -3,5714148 | 0,00035506 | 0,0168474 |
| Tnfrsf1a | 63,4539024 | -0,752567278 | 0,2095417 | -3,5703415 | 0,00035652 | 0,01686947 |
| Cybrd1 | 33,1987249 | -0,809360682 | 0,22792372 | -3,5631429 | 0,00036644 | 0,01729093 |
| Trim12c | 31,3949017 | -0,787610397 | 0,22272607 | -3,559883 | 0,00037102 | 0,01741027 |

| | | | | | | |
|---|---|---|---|---|---|---|
| Stxbp2 | 279,891188 | -0,705946369 | 0,19836181 | -3,5589294 | 0,00037237 | 0,01742546 |
| Gdf15 | 6,22527598 | -0,77657428 | 0,20292852 | -3,5542137 | 0,00037911 | 0,01764372 |
| Gabrb1 | 1791,24488 | 0,594208901 | 0,16737281 | 3,55172669 | 0,00038271 | 0,01766612 |
| Kctd4 | 75,9358672 | 0,727088946 | 0,20403977 | 3,55214431 | 0,00038211 | 0,01766612 |
| Tgfbr2 | 112,083614 | -0,707212327 | 0,19877586 | -3,5474606 | 0,00038896 | 0,01781338 |
| Kcnk12 | 62,4200526 | -0,730900148 | 0,20493422 | -3,5474035 | 0,00038905 | 0,01781338 |
| Lmo2 | 63,2969493 | -0,662962169 | 0,1861413 | -3,5461624 | 0,00039089 | 0,01784937 |
| Arhgap30 | 39,2646258 | -0,785404407 | 0,21968046 | -3,5392794 | 0,00040122 | 0,01827222 |
| Kcnj4 | 290,725852 | -0,60039235 | 0,17004975 | -3,5381846 | 0,00040289 | 0,0182991 |
| Ikbke | 20,4833671 | -0,759444768 | 0,22897738 | -3,5371373 | 0,00040449 | 0,01832283 |
| Zfp36 | 23,8976833 | -0,810810238 | 0,2271909 | -3,529764 | 0,00041593 | 0,01875645 |
| Dock8 | 82,8618672 | -0,661505711 | 0,1872986 | -3,529548 | 0,00041627 | 0,01875645 |
| B230334C09 | 1587,95853 | 0,616935799 | 0,1747581 | 3,52621417 | 0,00042155 | 0,01894392 |
| Cartpt | 9,36390099 | 0,627734234 | 0,20631734 | 3,52275384 | 0,00042709 | 0,01905679 |
| Trim26 | 121,156351 | -0,585951373 | 0,16633773 | -3,5238329 | 0,00042535 | 0,01905679 |
| Emp1 | 29,750626 | -0,792668617 | 0,22423437 | -3,5188059 | 0,00043349 | 0,01927685 |
| Gm28271 | 8,81115322 | -0,796594756 | 0,21904016 | -3,516171 | 0,00043782 | 0,01941835 |
| Gbp6 | 17,0608616 | -0,790344679 | 0,22854832 | -3,5137212 | 0,00044188 | 0,01954727 |
| Fblim1 | 10,4930946 | -0,756566357 | 0,21925721 | -3,5000191 | 0,00046522 | 0,02052669 |
| Tlcd3a | 127,781806 | 0,629340123 | 0,18025494 | 3,48865859 | 0,00048545 | 0,02125348 |
| Cxcl9 | 7,07161746 | -0,682509994 | 0,20079412 | -3,4835821 | 0,00049475 | 0,02151873 |
| Ms4a4b | 6,01995419 | -0,644339702 | 0,2055191 | -3,4832803 | 0,00049531 | 0,02151873 |
| 4933412O06 | 36,6192853 | 0,785716536 | 0,21898917 | 3,48022656 | 0,00050099 | 0,02170999 |
| Lbh | 202,220264 | -0,632404398 | 0,1818573 | -3,4740242 | 0,00051271 | 0,02204933 |
| Myo1f | 53,3918111 | -0,742394632 | 0,21361424 | -3,4719225 | 0,00051675 | 0,02216656 |
| Cst7 | 11,8375355 | -0,621232735 | 0,21141814 | -3,4666725 | 0,00052694 | 0,02243406 |
| S1pr3 | 52,6209001 | -0,715399956 | 0,20447108 | -3,4671974 | 0,00052592 | 0,02243406 |
| Jund | 543,644683 | -0,700763015 | 0,20102645 | -3,4652698 | 0,0005297 | 0,02243893 |
| H2-M3 | 17,8809903 | -0,783362347 | 0,2285747 | -3,4655323 | 0,00052918 | 0,02243893 |
| Cd84 | 53,0008717 | -0,753909501 | 0,21604087 | -3,4640983 | 0,00053201 | 0,02248024 |
| Piwil2 | 14,4911981 | -0,765587491 | 0,22833707 | -3,4558165 | 0,00054863 | 0,02295457 |
| Rps2 | 567,660881 | -0,584492164 | 0,16935229 | -3,4524337 | 0,00055555 | 0,02313799 |
| Tgif1 | 16,0721254 | -0,789623875 | 0,22858463 | -3,452345 | 0,00055574 | 0,02313799 |
| Ptpn6 | 32,0820625 | -0,76831254 | 0,22311667 | -3,4488909 | 0,00056289 | 0,02333719 |
| Il2rb | 9,53396094 | -0,602512981 | 0,206276 | -3,4454273 | 0,00057016 | 0,02356511 |
| Tmem140 | 18,9280154 | -0,729850189 | 0,22798236 | -3,4417355 | 0,000578 | 0,02371595 |
| Trpc6 | 74,9967221 | 0,663133232 | 0,19164075 | 3,4419045 | 0,00057763 | 0,02371595 |
| Plaat1 | 63,883824 | 0,664339591 | 0,19295053 | 3,44253116 | 0,0005763 | 0,02371595 |
| S100a6 | 19,6817092 | -0,812737013 | 0,22837741 | -3,4390985 | 0,00058365 | 0,02389047 |
| Il10ra | 48,9666738 | -0,686888664 | 0,19995754 | -3,4342868 | 0,00059412 | 0,0241441 |
| Rab15 | 794,241334 | -0,597955583 | 0,17422332 | -3,4297482 | 0,00060414 | 0,02443465 |
| Gm42517 | 249,247804 | -0,600391248 | 0,17501024 | -3,4276494 | 0,00060883 | 0,02456583 |
| Hk2 | 63,6681626 | -0,715734186 | 0,20950202 | -3,4237888 | 0,00061755 | 0,02479965 |
| Phf1 | 309,151213 | -0,685573729 | 0,20056568 | -3,4202836 | 0,00062556 | 0,02506219 |
| Ppp1r35 | 27,4854786 | -0,788561274 | 0,22855548 | -3,4140029 | 0,00064016 | 0,02558678 |
| Ackr3 | 44,091602 | 0,738554848 | 0,21579372 | 3,40823468 | 0,00065385 | 0,02607251 |

| | | | | | | |
|---|---|---|---|---|---|---|
| Tgfb1i1 | 61,7467103 | -0,670460828 | 0,19597985 | -3,4054656 | 0,00066051 | 0,02616337 |
| Crybg1 | 14,5750201 | -0,755510005 | 0,2262143 | -3,4053696 | 0,00066075 | 0,02616337 |
| Clec7a | 30,9303158 | -0,794948184 | 0,2277517 | -3,4019145 | 0,00066916 | 0,02643477 |
| Stat6 | 77,8287876 | -0,612602269 | 0,18065755 | -3,3809779 | 0,00072228 | 0,02827058 |
| Cd63 | 94,1928476 | -0,719684765 | 0,21184783 | -3,3816883 | 0,00072042 | 0,02827058 |
| Trim47 | 21,6769924 | -0,7876343 | 0,22878576 | -3,3828098 | 0,00071748 | 0,02827058 |
| Gas6 | 342,975209 | -0,586824307 | 0,17381494 | -3,3798677 | 0,00072521 | 0,02831976 |
| Psrc1 | 49,4424331 | -0,68826992 | 0,20135341 | -3,3774704 | 0,00073156 | 0,02850225 |
| Lamb1 | 220,535551 | 0,605544338 | 0,17936201 | 3,37589961 | 0,00073575 | 0,02859987 |
| Tcf24 | 6,96911004 | -0,691536271 | 0,21140837 | -3,3704869 | 0,00075035 | 0,0290348 |
| Tns1 | 320,815797 | -0,587988676 | 0,1743005 | -3,3689447 | 0,00075457 | 0,02913138 |
| Cfap54 | 68,710081 | 0,668913055 | 0,19813184 | 3,36825192 | 0,00075646 | 0,02913846 |
| Smim17 | 66,0591606 | 0,650190702 | 0,1912948 | 3,3671998 | 0,00075936 | 0,02918369 |
| Slc11a1 | 29,9233692 | -0,74561484 | 0,22161487 | -3,3651749 | 0,00076495 | 0,02926629 |
| Cdk2ap1 | 44,5897157 | -0,677400198 | 0,20027301 | -3,3626068 | 0,0007721 | 0,0294735 |
| Bend5 | 63,4911166 | -0,634011988 | 0,18900685 | -3,3585651 | 0,00078348 | 0,02984085 |
| Phf11b | 6,04545685 | -0,682216806 | 0,19797158 | -3,357921 | 0,00078531 | 0,02984356 |
| Notch3 | 128,761461 | -0,664722257 | 0,19703854 | -3,3555433 | 0,00079209 | 0,0300341 |
| Fam181b | 66,0460227 | -0,742887974 | 0,21902196 | -3,3532185 | 0,00079878 | 0,03022007 |
| Smim13 | 1062,69697 | 0,605913636 | 0,18057137 | 3,35189086 | 0,00080262 | 0,03029788 |
| Cwh43 | 23,3270577 | -0,755684858 | 0,22404304 | -3,347442 | 0,00081561 | 0,03061872 |
| Tgtp2 | 13,6756442 | -0,652150524 | 0,22132282 | -3,347132 | 0,00081652 | 0,03061872 |
| Gimap3 | 9,29512008 | -0,692847611 | 0,21933599 | -3,3447605 | 0,00082354 | 0,03081369 |
| Gm32856 | 32,0486354 | 0,718670888 | 0,21332734 | 3,33699094 | 0,00084691 | 0,03154915 |
| Fam174a | 188,173754 | -0,596880261 | 0,17944892 | -3,3305861 | 0,00086663 | 0,03206546 |
| Thbs1 | 38,2160132 | -0,744697491 | 0,22869548 | -3,3269472 | 0,00087803 | 0,03235379 |
| Vasp | 66,0339557 | -0,67977726 | 0,20447315 | -3,3254285 | 0,00088283 | 0,03246015 |
| Cd52 | 12,1005257 | -0,675934046 | 0,21767495 | -3,3227252 | 0,00089143 | 0,03260207 |
| Atp2b4 | 215,37359 | -0,633506106 | 0,19092539 | -3,3070861 | 0,00094272 | 0,03421788 |
| Psme1 | 84,2260115 | -0,631965826 | 0,19083086 | -3,3042491 | 0,00095231 | 0,03449237 |
| Mical1 | 51,3575662 | -0,641465848 | 0,19386924 | -3,3021075 | 0,00095961 | 0,03468286 |
| Penk | 268,585283 | -0,703000991 | 0,21218567 | -3,2970615 | 0,00097702 | 0,03516241 |
| Col1a1 | 33,3218408 | -0,789944877 | 0,22869916 | -3,2971238 | 0,0009768 | 0,03516241 |
| Plscr4 | 62,3002844 | 0,68154907 | 0,20389612 | 3,28335216 | 0,0010258 | 0,03676232 |
| Vamp8 | 18,1227066 | -0,766962147 | 0,22871836 | -3,2784133 | 0,00104392 | 0,03708526 |
| Dera | 18,381944 | -0,75687066 | 0,22836415 | -3,2765601 | 0,0010508 | 0,03711004 |
| Glt8d2 | 204,394826 | 0,604136164 | 0,18356993 | 3,27369182 | 0,00106152 | 0,03733353 |
| Cd180 | 31,9725406 | -0,749416397 | 0,22789485 | -3,2723041 | 0,00106675 | 0,03736256 |
| Aif1 | 19,1314452 | -0,754157014 | 0,22787988 | -3,2728237 | 0,00106479 | 0,03736256 |
| Phlda1 | 54,490484 | -0,645951591 | 0,19600098 | -3,2716932 | 0,00106906 | 0,03736634 |
| Aff2 | 64,8417264 | 0,634041132 | 0,19369022 | 3,26477532 | 0,00109551 | 0,03821236 |
| Ncf1 | 48,7910791 | -0,676937455 | 0,20957712 | -3,2625109 | 0,0011043 | 0,03844003 |
| Gm14443 | 112,421037 | 0,670194209 | 0,2033195 | 3,26045472 | 0,00111234 | 0,03864064 |
| Mpeg1 | 264,394916 | -0,744130552 | 0,22884725 | -3,25853 | 0,00111991 | 0,03882433 |
| Lpcat2 | 80,7126349 | -0,585700696 | 0,17933904 | -3,2549687 | 0,00113405 | 0,03911135 |
| Csf2rb | 22,1709742 | -0,74957114 | 0,22730326 | -3,2462037 | 0,00116955 | 0,04003756 |

| | | | | | | |
|---|---|---|---|---|---|---|
| Adap2 | 52,2186404 | -0,660172545 | 0,20511612 | -3,2417559 | 0,00118796 | 0,04044053 |
| Vwa1 | 82,4986526 | -0,645178243 | 0,19784453 | -3,2379026 | 0,00120412 | 0,04082709 |
| Spock1 | 658,133627 | 0,597836788 | 0,18522146 | 3,23666066 | 0,00120937 | 0,04092349 |
| Shisa8 | 13,9237841 | -0,759886599 | 0,22733129 | -3,2308851 | 0,00123408 | 0,04159373 |
| Slc39a1 | 198,631522 | -0,593042388 | 0,18331019 | -3,2262521 | 0,00125423 | 0,04202284 |
| Tnfrsf1b | 36,542043 | -0,705323001 | 0,21913194 | -3,226501 | 0,00125314 | 0,04202284 |
| Sertad1 | 28,3059163 | -0,719393069 | 0,22144356 | -3,2271635 | 0,00125024 | 0,04202284 |
| Tgfb1 | 28,3178216 | -0,699559083 | 0,22185307 | -3,2218557 | 0,00127363 | 0,04226948 |
| Hmox1 | 24,7362951 | -0,701261843 | 0,22449095 | -3,2222679 | 0,0012718 | 0,04226948 |
| Cd68 | 32,1336278 | -0,722444143 | 0,22811647 | -3,221766 | 0,00127403 | 0,04226948 |
| Crhbp | 129,222571 | 0,646618762 | 0,20018982 | 3,22242219 | 0,00127112 | 0,04226948 |
| Dact3 | 774,103713 | -0,636357795 | 0,19640859 | -3,2200033 | 0,00128189 | 0,04241018 |
| Rbl1 | 16,6554676 | -0,744591613 | 0,22819988 | -3,2181429 | 0,00129024 | 0,04247525 |
| Ccdc88b | 33,8018155 | -0,701744798 | 0,22884026 | -3,2181792 | 0,00129007 | 0,04247525 |
| Gm13067 | 8,11469683 | -0,715462783 | 0,21803777 | -3,2155834 | 0,0013018 | 0,04277295 |
| Fgf10 | 120,061709 | 0,590536177 | 0,18335746 | 3,20932164 | 0,00133049 | 0,04346338 |
| Rrbp1 | 171,935836 | -0,635132175 | 0,19929543 | -3,1979293 | 0,00138418 | 0,04470171 |
| Socs3 | 14,4113254 | -0,763068472 | 0,22877472 | -3,1928482 | 0,00140877 | 0,04514483 |
| Tgfb3 | 74,8063118 | -0,604568777 | 0,19013017 | -3,1923522 | 0,00141119 | 0,04514483 |
| Shroom2 | 445,542806 | 0,58539547 | 0,18316551 | 3,19392962 | 0,0014035 | 0,04514483 |
| Pdrg1 | 81,1957275 | -0,584730628 | 0,18360522 | -3,1897986 | 0,00142372 | 0,04546001 |
| Gcnt2 | 124,107134 | -0,612811761 | 0,19177799 | -3,1877937 | 0,00143363 | 0,04569049 |
| Tcf15 | 5,02987916 | -0,63499808 | 0,19405801 | -3,1847623 | 0,00144873 | 0,04608533 |
| Lgals3 | 8,15347253 | -0,670470017 | 0,21465591 | -3,1838589 | 0,00145326 | 0,04614299 |
| Degs1 | 168,449579 | 0,627142022 | 0,19875242 | 3,16831609 | 0,00153325 | 0,04796551 |
| Mir341 | 42,1209292 | 0,723945068 | 0,22656292 | 3,16578979 | 0,00154663 | 0,04820646 |
| Rasal3 | 51,8521439 | -0,684977646 | 0,2186752 | -3,165859 | 0,00154626 | 0,04820646 |
| Tlr9 | 13,8332931 | -0,709065505 | 0,22887425 | -3,1578434 | 0,00158941 | 0,04917905 |
| Nudt17 | 28,5869538 | 0,690142649 | 0,21733444 | 3,15706378 | 0,00159366 | 0,04922106 |
| Arhgap11a | 25,403789 | -0,706503955 | 0,22479351 | -3,1548263 | 0,00160594 | 0,04924197 |
| Renbp | 12,78096 | -0,701844805 | 0,21784254 | -3,1549518 | 0,00160525 | 0,04924197 |
| Gm37661 | 50,3367076 | 0,612963733 | 0,19450046 | 3,15391052 | 0,00161098 | 0,04930774 |
| Glipr2 | 14,2056493 | -0,71669229 | 0,22805054 | -3,1518297 | 0,00162251 | 0,04948214 |
| Sf3b5 | 80,0209131 | -0,68414719 | 0,21719069 | -3,1522774 | 0,00162002 | 0,04948214 |
| Lyn | 63,1363707 | -0,679813517 | 0,21531649 | -3,1486038 | 0,00164052 | 0,04976356 |

**Supplementary Table 6 -** Table summarizing the effect of normalization with DESeq2

| Sample ID | Phenotype | Total raw counts | Total normalized counts |
|---|---|---|---|
| A1 | scramble | 6432443 | 5281271 |
| A2 | scramble | 4367843 | 5339928 |
| A3 | scramble | 6899426 | 5243021 |
| B1 | shRNF10 | 6537932 | 5373401 |
| B2 | shRNF10 | 5767767 | 5497725 |
| B3 | shRNF10 | 5352238 | 5393735 |
